# Hydrophobic mismatch drives self-organization of designer proteins into synthetic membranes

**DOI:** 10.1101/2022.06.01.494374

**Authors:** Justin A. Peruzzi, Jan Steinkühler, Timothy Q. Vu, Taylor F. Gunnels, Peilong Lu, David Baker, Neha P. Kamat

## Abstract

The extent to which membrane biophysical properties, such as hydrophobic thickness, can drive membrane protein organization remains unknown. Inspired by this question, we used *de novo* protein design, molecular dynamic simulations, and cell-free systems to elucidate how membrane-protein hydrophobic mismatch affects protein integration and organization in synthetic lipid membranes. We found that membranes must deform to accommodate membrane-protein hydrophobic mismatch, which reduces the expression and co-translational insertion of membrane proteins into synthetic membranes. We used this principle to sort proteins both between and within membranes, thereby achieving one-pot assembly of vesicles with distinct functions and controlled split-protein assembly, respectively. Our results shed light on protein organization in biological membranes and provide a framework to self-organizing membrane-based materials with new functions.

## Main Text

Biological cells leverage membrane bound compartments to perform complex functions with precise spatial and temporal control. To execute these processes, cells must insert and sort proteins into distinct membrane compartments. Cellular membranes possess a variety of mechanisms to control membrane protein location. Protein transport is largely mediated by different protein-protein interactions and protein machinery such as clathrin, COPI, and SNARE proteins (*1, 2*). However, it has also been hypothesized that lipid-protein interactions can drive inter- and intramembrane protein organization (*3*–*5*). Membranes and membrane-proteins have been shown to possess complementary physiochemical properties, likely allowing for proper protein sorting and function within cells (*5*–*7*). Specifically, protein transmembrane domain length and geometry have been shown to correlate with protein localization between and within different cellular membranes (*6, 7*). These studies suggest that physical features of membranes and proteins, such as the hydrophobic thickness of transmembrane domains and lipid bilayers, are used by cells to organize membrane proteins into distinct organelle membranes, thereby controlling membrane-based behaviors.

A major challenge to probing the contribution of physical factors in membrane protein folding and sorting is the complexity of biological cells. Cells possess a diverse lipidome, unique protein structures, and complex protein sorting machinery, making it difficult to parse out the extent to which specific lipid-protein biophysical interactions influence membrane protein folding and trafficking (*2, 8*). Recent advances in *de novo* protein design and membrane-augmented cell-free protein synthesis systems offer a route to explore how protein and lipid properties affect membrane protein integration and dynamics in a controlled environment.

Developing *in vitro* methods to characterize specific membrane-protein interactions, such as those influenced by membrane physical properties or protein sequence and structure, will shed light on fundamental biological questions surrounding protein folding and sorting. In addition, this insight will enable the design of membrane-based materials (e.g. biosensors, drug delivery vehicles) with properties beyond what is possible in nature (*9*–*14*), critical in advancing applications in biosensing and therapeutics.

Towards this goal, we designed alpha-helical hairpin dimers ranging from 10 to 50 Å in hydrophobic length, and transmembrane pores ranging from 20 to 50 Å in hydrophobic length based on previous protein design scaffolds (*15, 16*), and assessed these proteins’ ability to insert and organize into a thin (14:1 PC, 23 Å), medium (18:1 PC, 29 Å), and thick lipid membranes (22:1 PC, 37 Å)(*17*). Together with molecular dynamic (MD) simulations, and experimental studies using cell-free systems, we were able to systematically probe and characterize how membrane-protein hydrophobic mismatch, affects protein expression, co-translational folding, and location within a membrane in a way that was not possible to date.

### Hydrophobic mismatch reduces co-translational insertion of designed proteins

We first assessed how hydrophobic mismatch, defined here as membrane hydrophobic thickness minus the protein hydrophobic thickness, affects co-translational insertion of *de novo* designed hairpin proteins (Fig. 1A). We performed MD simulations of proteins in a thin, medium, and thick lipid system and measured membrane thickness as a function of distance from the protein and found that membranes deform close to the protein insertion site, deforming more as hydrophobic mismatch increases (Fig. 1B, 1C, S1). We then experimentally characterized how hydrophobic mismatch impacted protein expression *in vitro*. We designed plasmids encoding hairpin proteins. A C-terminal monomeric-enhanced GFP allowed us to monitor expression and proper folding of proteins by GFP fluorescence (*18, 19*). By adding a plasmid encoding a membrane protein and pre-assembled phospholipid vesicles to a cell-free protein synthesis system, we could track expression and cotranslational insertion of designed proteins into synthetic membranes of the vesicles (Fig. 1D, S2). Proteins were expressed in the presence of no membrane or a thin, medium, or thick lipid membrane to match our simulations. We monitored protein folding via GFP fluorescence and measured protein expression via western blots. All designed proteins expressed poorly in the absence of vesicles, suggesting that their enhanced expression in the presence of membranes is due to co-translational protein folding and insertion into synthetic membranes (Fig. 1E, S3). When comparing the expression of each protein in the presence of the three lipid systems, we found that expression and proper folding was generally maximized when membrane-protein hydrophobic mismatch was minimized for each studied protein (Fig. 1E, Fig. S3, S4). Furthermore, GFP fluorescence, determined experimentally, correlated linearly with membrane compression, determined computationally (Fig. 1F). Together, these data demonstrate that membrane-protein hydrophobic mismatch inhibits membrane protein expression.

**Figure 1.**
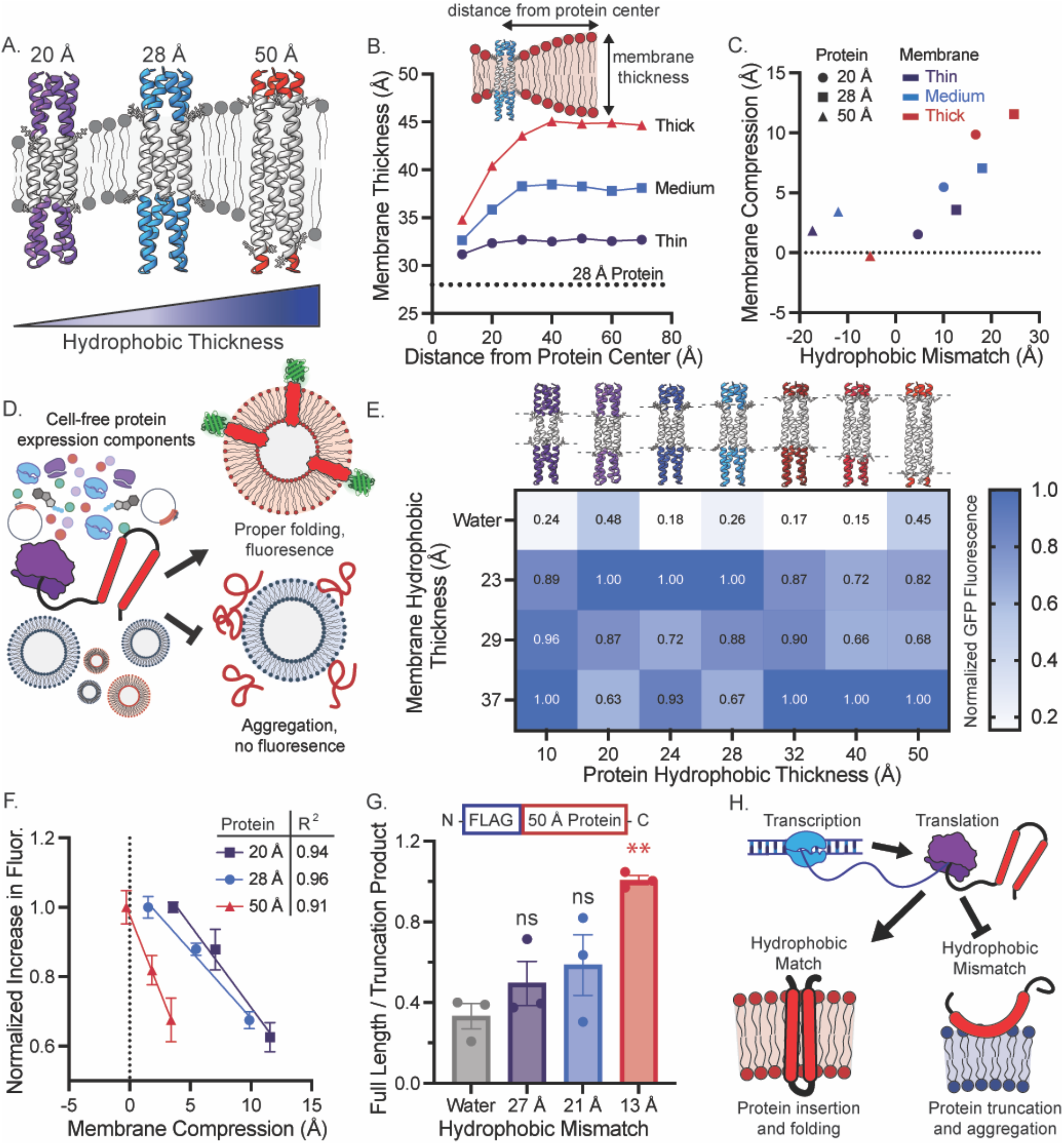
Minimizing hydrophobic mismatch maximizes cell-free expression of membrane proteins into synthetic membranes. (**A**) Interactions between *de novo* designed membrane proteins of varying hydrophobic thicknesses and synthetic membranes were explored. (**B**) Computationally probing membrane thickness as a function of distance from an inserted, 28 Å protein reveals that membranes must deform more to accommodate larger hydrophobic mismatch. The horizontal line represents the protein hydrophobic thickness. (**C**) Membrane compression is positively correlated with hydrophobic mismatch. Datapoints derived from simulations of the 20, 28, and 50 Å proteins for the three membrane compositions (thin, medium, thick). (**D**) The effect of hydrophobic mismatch on protein expression and folding in a cell-free protein synthesis systems was explored using mEGFP as a folding reporter. (**E**) Protein expression, as measured by increased GFP fluorescence, is maximized in hydrophobically matched membranes. Values represent the mean of 3 independent replicates, normalized to the maximum increase in fluorescence for each protein construct. (**F**) Increase in GFP fluorescence is linearly correlated with membrane compression, as measured by MD simulation. (**G**) The proportion of full length 50 Å protein relative to incomplete proteins, increases as hydrophobic mismatch is minimized. **p < 0.001, nonsignificant (ns). P-values were generated by a one-way ANOVA using Dunnett’s multiple comparison test, comparing everything to the water sample. All error bars represent the S. E. M. for *n=3* independent replicates. (**H**) Schematic of cell-free protein expression of proteins into membranes of different hydrophobic thicknesses. Proteins insert and fold best into hydrophobically matched membranes.

We next examined how transcription and translation were affected by hydrophobic mismatch. We measured transcription by adding the DNA sequence for the malachite green aptamer immediately after the gene encoding the 50 Å protein. As the aptamer is transcribed, it binds to malachite green and dye fluorescence increases (*20*). The presence of vesicles was found to inhibit transcription of the 50 Å protein; however, no significant differences in malachite green fluorescence were observed between the three lipid systems suggesting hydrophobic mismatch does not measurably affect transcription (Fig. S5)(*18*). To characterize how translation is affected by membrane-protein hydrophobic mismatch, we added a N-terminal FLAG tag to the 50 Å protein, allowing us to monitor the formation of both full-length and truncated protein products. Analyzing the expression of 50 Å by western blot, we observed an increase in incomplete protein products relative to full-length protein as a function of hydrophobic mismatch (Fig. 1G, S6). The higher proportion of truncated protein products in hydrophobically mismatched systems suggests that translation is affected by hydrophobic mismatch. We hypothesize this effect arises from the energy cost of deforming membranes to accommodate differences in hydrophobic mismatch, which reduces the probability of protein co-translational insertion and proper folding. Misfolded proteins likely capture nascent proteins from the ribosome, thus reducing the rate of protein insertion and folding, and increasing the frequency of incomplete translation (Fig. 1H).

### Protein-lipid hydrophobic matching can be used to organize proteins between membranes and impart differentiated functionality

The differential expression and integration of membrane proteins into membranes of different thicknesses raised the possibility that this physical phenomenon could be used to enrich select populations of vesicles with a membrane protein in one pot. Based on the designs of our previous work, we created transmembrane pore proteins with a constitutively open 10 Å pore and with hydrophobic thicknesses ranging from 20 to 50 Å (Fig. 2A) (*16*). To validate that pore proteins inserted into membranes, we expressed them in the presence of vesicles encapsulating calcein, a self-quenching dye (Fig. 2B). Upon expression of pore proteins, we observed calcein leakage and increased fluorescence (*21*). We first confirmed that calcein leakage was specific to pore insertion (Fig. S7), and then expressed the pores in the presence of vesicles with thick or thin membranes, encapsulating calcein. When normalized by protein expression, as determined by western blot, hydrophobically matched proteins released the most amount of calcein (Fig. 2C, D). This result demonstrates that reducing hydrophobic mismatch between the designed pore proteins and membranes maximizes the functional incorporation of membrane proteins.

**Figure 2.**
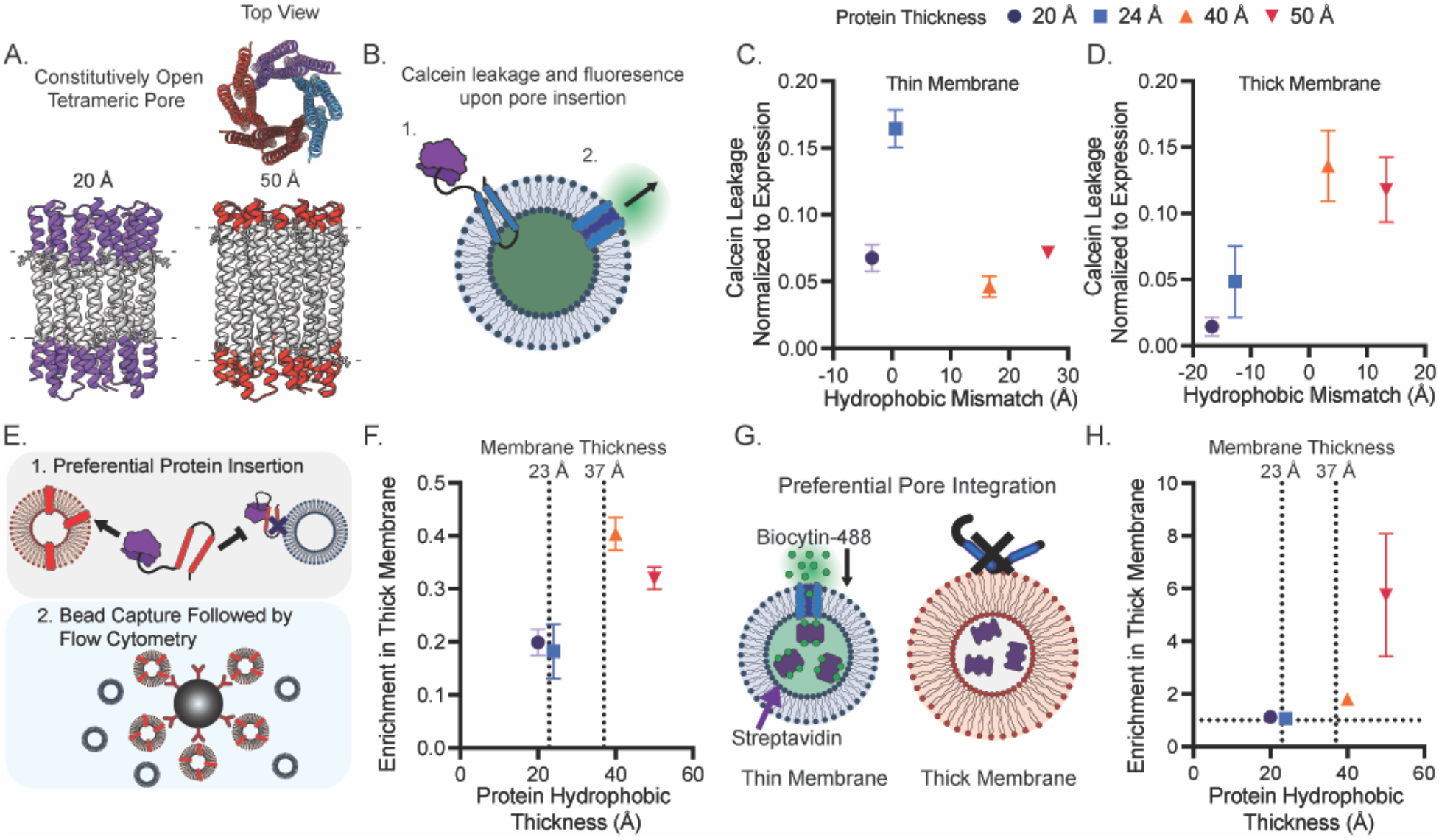
Hydrophobic mismatch alone can organize cell-free expressed proteins between distinct membrane compartments. (**A**) Constitutively open pore proteins of varying hydrophobic thicknesses were designed. (**B**) Proper folding and insertion of pore proteins was assessed via calcein leakage. Calcein leakage through *de novo* designed channel proteins is maximized when hydrophobic mismatch is minimized in both thin (**C**) and thick (**D**) membranes. Expression of protein in the presence of two populations of vesicles, followed by bead capture and flow cytometry enable the characterization of protein organization **(E)**. As hydrophobic thickness of designed membrane channels is increased, the protein-mediated binding of thick membrane vesicles to beads increases relative to thin membrane vesicles (**F**). (**G**) To selectively deliver cargo in a mixed vesicle population, we expressed pore proteins in the presence of thick and thin membranes, encapsulating streptavidin. Following protein expression, vesicles were incubated with AF488-biocytin, which could enter vesicles following pore integration. The amount of dye that was delivered to each population of vesicles was measured by flow cytometry. Vertical dotted lines in (**F**) and (**H**) correspond to membrane hydrophobic thickness. (**H**) Cargo delivery to thick and thin vesicles could be tuned by hydrophobic thickness of designed membrane pores. All experiments were performed 3 times, error bars represent the S. E. M.

We next assessed the extent of protein expression and folding of a single protein (20, 24, 40, or 50 Å in hydrophobic thickness) into thick and thin membranes (37 and 23 Å respectively), when both membranes were present within one reaction. To evaluate differential integration, we developed a flow cytometry-based assay where each set of vesicles was labeled with an orthogonal lipid conjugated dye and each protein contained a C-terminal FLAG tag. Proteins were expressed in the presence of vesicles and were collected with anti-FLAG antibody-conjugated beads. Beads were analyzed by flow cytometry and read for colocalized vesicle fluorescence, which should occur by way of interactions of membrane-integrated proteins with the beads (Fig. 2E). We monitored the ratio of fluorescence from the thick membranes to the thin membranes that was colocalized to the beads and found that as hydrophobic thickness of a protein increased, this ratio increased. This result indicates that proteins preferentially fold into hydrophobically matched membranes (Fig. 2F, S8) and that protein enrichment in one population of membranes can be increased relative to another vesicle population by minimizing hydrophobic mismatch.

We explored the capacity of hydrophobic mismatch to assemble vesicles with a distinct functionality, in this case enhanced permeability, due to preferred integration of membrane proteins. We measured membrane permeability to a biotinylated fluorophore (∼1 kDa)(*16*). We encapsulated streptavidin in the lumen of thick and thin membranes, each labeled with a distinct lipid-conjugated fluorescent dye. We expressed proteins of different hydrophobic thicknesses in the presence of both vesicles, purified away free streptavidin, and then incubated the vesicles with biocytin conjugated AlexaFluor 488. Biocytin entry into vesicles, which should vary as a function of the number of functional pores in each vesicle membrane, could be monitored via vesicle-localized biocytin fluorescence since biocytin cannot leave the vesicles after it is bound to streptavidin in the vesicle lumen (Fig. 2G). Samples were then analyzed via flow cytometry and the ratio of thick and thin vesicles encapsulating AlexaFluor 488 were compared. As protein hydrophobic thickness increased, we observed an increase in this ratio suggesting that the population of vesicles to which biotin-AlexaFluor 488 was preferentially delivered could be tuned by biasing protein integration into hydrophobically matched vesicles membranes (Fig. 2H, S9). Together, these data suggest that membrane compartments can be enriched with distinct protein content and therefore endowed with distinct function by modulating lipid-protein hydrophobic mismatch. Our results highlight the capacity of protein-membrane hydrophobic mismatch alone to organize proteins between distinct membranes *in vitro* and suggest a route to engineer more complex membrane-based materials, such as differentiated-nested vesicles or synthetic organelles.

### Hydrophobic mismatch coupled with phase separating lipid mixtures controls protein-protein interactions within a single membrane

The lateral organization of membrane proteins in a single membrane is important to control protein-protein and protein-lipid interactions and subsequent signaling activity (*7, 22*). This organization arises due to different lipid-lipid, lipid-protein, and cytoskeletal interactions. While the functional relationship between protein organization and signaling has been explored in cellular contexts, it has not yet been recapitulated *in vitro*. Demonstrating this organization experimentally is critical to identify the molecular and physical origin of these interactions. In doing so, we not only uncover the extent to which protein and lipid driven organization may enable protein organization in cells, but also provide a route to design more complex sensing and signaling modalities within membrane-based materials.

Previous work has demonstrated that peptides can be laterally organized through membrane ordering in unsaturated lipid systems (*23*) and that beta-barrel proteins can associate with liquid-ordered lipid phases through the modulation of protein hydrophobic thickness (*24*). However, contradicting phase behavior of proteins in cellular, *in silico*, and synthetic membranes has been noted (*25, 26*), likely due to the use of microdomain forming lipid mixtures in synthetic lipid systems, which are more ordered than biological membranes, hindering protein association with ordered lipid phases. We hypothesize that by designing membranes just above a lipid de-mixing transition, like biological membranes (*27*), we could observe induction of lipid domains induced by local changes in curvature or hydrophobic thickness around membrane components, such as proteins (*28*–*30*).

To examine the ability of hydrophobic mismatch to affect protein location and protein-protein interactions in a single membrane, we first characterized how single proteins co-localize with lipid components based on hydrophobic mismatch. We prepared membranes with a shorter unsaturated lipid, 14:1 PC, and a thicker saturated lipid, DPPC (16:0 PC), and cholesterol. This combination of lipids is prone to phase separation and at different lipid ratios can form homogenous and phase separated membranes (*31*). We chose a lipid composition just above a de-mixing transition (*25, 26, 32*) (Fig. S10). We simulated membrane interactions with thin (20 Å) and thick (50 Å) proteins using coarse-grained MD simulations of lipid composition comparable to the experimental system. We observed that insertion of membrane proteins into an initially homogenous lipid mixture induced lipid reorganization. Distinct lipid-protein domains formed: a domain rich in thinner, unsaturated lipid (DyPC) appeared around the 20 Å protein and a domain rich in the thicker, saturated lipid (DPPC) appeared around the 50 Å protein over time (Fig. 3A). We then studied the effect of increasing temperature, a means to dissolve lipid-protein domains as saturated and unsaturated lipids become more miscible at elevated temperatures (*31*). As temperature increased, protein-lipid contacts converged toward the average composition of the membrane.

**Figure 3.**
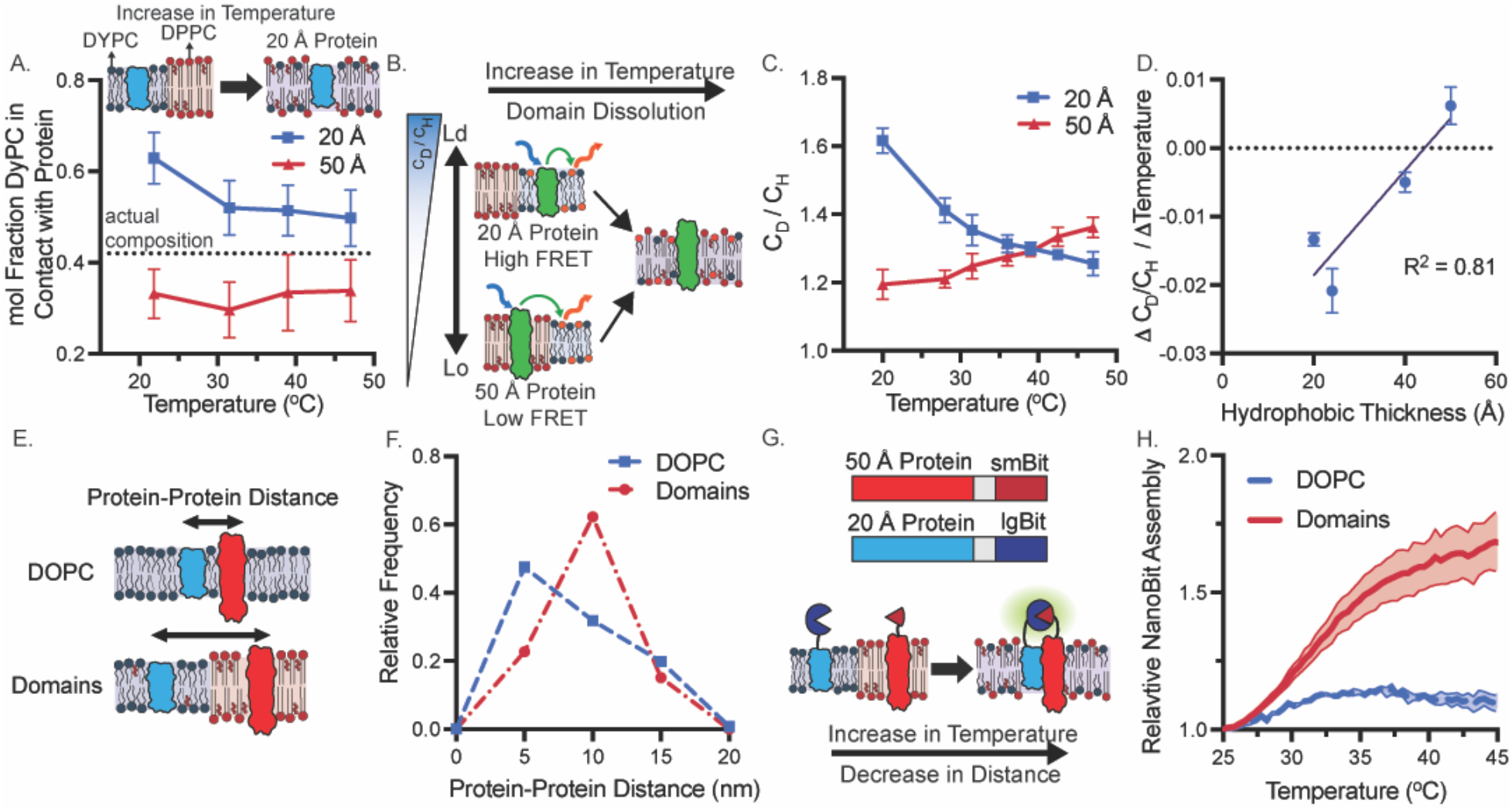
Lipid-protein hydrophobic mismatch can dynamically tune protein-protein interactions. (**A**) MD simulations indicate the 20 Å hairpin proteins interact more with the shorter lipid, DyPC, compared to the 50 Å protein. As temperature increases, the protein-DyPC contacts shift towards the average membrane composition. The dotted line indicates the actual composition of DyPC, 42.5 mol%. (**B**) Lipid-protein FRET between C-terminal AlexaFluor 488-SNAP tag and Rhodamine conjugated lipids enable the evaluation of protein organization within synthetic membranes. (**C**) 20 Å and 50 Å proteins associate differently with Rhodamine conjugated lipids, as reported by C_D_/C_H_. As temperature increases, the membrane becomes more fluid enabling lipid mixing and convergence of the two C_D_/C_H_ curves. (**D**) Total change in C_D_/C_H_ from 20 to 45°C correlates with protein hydrophobic thickness. (**E, F**) Protein-protein distance can be modulated by lipid composition of synthetic membranes. In homogenous DOPC membranes, 20 Å and 50 Å proteins can be close to one another, however in phase separating lipid mixtures proteins remain farther from one another as predicted by MD simulations. (**G, H**) Lipid domain forming membranes compartmentalize and separate 20 Å and 50 Å proteins at room temperature. Upon heating, to enable protein and lipid mixing, split luciferase reconstitution and subsequent luminescence is higher in the domain forming lipid mixture, compared to DOPC. All experiments were performed 3 times, error bars represent the S. E. M.

We then experimentally assessed how proteins were organized in our membranes via lipid-protein FRET (fluorescence resonance energy transfer). To accomplish this, we added rhodamine dye conjugated to 18:1 PC into our membranes, which localizes with shorter, unsaturated lipids, and fused a C-terminal SNAP tag to each protein, allowing conjugation of AlexaFluor 488. Using FRET between SNAP Alexa Fluor 488 and the lipid-conjugated rhodamine dye, we could calculate the local concentration of rhodamine around the protein in domain forming membranes (14:1 PC/DPPC/Chol) compared to homogenous membranes (DOPC), represented as C_D_/C_H_ (Eq. 1). Using this metric, C_D_/C_H_ will be higher when proteins and dye partition to the same lipid domain and low when they partition to separate domains (fig. S11, Fig. 3B, fig. S12)(*24*). We measured C_D_/C_H_ values of the 20 and 50 Å protein over a range of temperatures. At room temperature, the 20 Å protein had a higher C_D_/C_H_, indicating that it resides in the dye containing, thinner and more unsaturated 14:1 PC rich phase. Upon increasing temperature to dissolve domains, we observed that the average distance between the unsaturated lipid dye and thinner protein increased. Conversely, the 50 Å protein had lower C_D_/C_H_ values at lower temperatures that increased with increasing temperature. This increase in C_D_/C_H_ values suggests the larger 50 Å protein shifted from an original position in the dye poor, thicker and more saturated DPPC rich lipid phase to one more well mixed with the lipid dye (Fig. 3C). Excitingly, the FRET data obtained as a function of temperature mirrors our simulation data in Fig. 3A quite well. Further, C_D_/C_H_ and its change over temperature for the 20, 24, 40, and 50 Å hairpin proteins correlate with hydrophobic thickness (Fig. 3D, fig. S12) and demonstrates lipid composition around a protein can be tuned by hydrophobic mismatch. Together the simulations and FRET data suggest that that hydrophobic mismatch coupled with lipid domain formation can be leveraged to organize proteins into distinct regions within synthetic membranes.

As a next step, we wondered if we could modulate protein-protein interactions by localizing proteins to separate lipid domains. Controlling the interactions between membrane proteins within a membrane would offer substantial advantages in the design of membrane-based technologies such as those that utilize transmembrane signaling transduction modules (*9, 10*). We first investigated how two proteins interact with one another by performing MD simulations of the 20 Å and 50 Å hairpin protein in a homogenous, single component and heterogenous, ternary membranes (Fig. 3E). These simulations demonstrated that proteins were able to be close to one another in homogenous membranes but remained separated in our mixed, phase segregated membrane (Fig. 3F, S13, Movie 1, S14).

We then investigated if this behavior could be recapitulated experimentally and used to control assembly of a split protein. We fused rapamycin inducible-dimerizing domains and NanoBit, a split nano luciferase (*33*), to the C terminus of the 20 and 50 Å proteins. To assess how lipid domains affected protein compartmentalization and subsequent NanoBit assembly, we co-expressed smBit and lgBit fused to the 20 Å and 50 Å protein, respectively, into homogenous or phase separated membranes. We observed how luminescence changed following the addition of rapamycin (*9*), which chemically induces protein dimerization and subsequent luciferase assembly. Upon addition of rapamycin, we saw an increase in luminescence when proteins were in homogenous membranes; however, we observed minimal increases in phase separated membranes, suggesting that proteins were unable to dimerize due to segregation into different lipid domains (Fig. S15). We then performed a temperature ramp on these systems to dissolve lipid domains (Fig. 3G). In phase separated systems, we observed an increase in luminescence with temperature when lgBit and smBit are fused to the hetero-pair of 20 and 50 Å protein, respectively, relative to when smBit and lgBit are both conjugated to homo-pairs of either the 20 or 50 Å proteins. Importantly, only a slight increase in luminescence is observed with increase in temperature when proteins are in homogenous membranes, indicating the proteins are more evenly distributed in the homogenous membrane (Fig. 3H, S16). Combined, the simulation and experimental data suggest lipid-lipid and lipid-protein interactions can together be harnessed to modulate protein interactions in a single membrane. Such control should improve the specificity and off target, ligand-independent activation of engineered receptor systems in both cellular and synthetic membranes (*14*).

## Conclusions

Here, we report on the capacity of hydrophobic mismatch between membranes and membrane proteins to drive changes in membrane protein synthesis and protein location. Using cell-free systems to recapitulate membrane protein synthesis and folding outside of the cellular environment, we demonstrate that not only do proteins insert and fold better into hydrophobically matched membranes, but that increases in mismatch reduce the yield of protein that is synthesized and increase the amount of incomplete, truncated translation products. Once in a mixed composition membrane capable of phase segregation, hydrophobic mismatch between lipids and proteins can drive reorganization to segregate lipids and proteins of similar length. Capitalizing on these inter-membrane and intra-membrane sorting mechanisms, we demonstrate: (1) the one-pot assembly of membranes with distinct protein incorporation and corresponding function based on hydrophobic mismatch alone and (2) control over the interactions of chemically-inducible dimerizing proteins in a single membrane. These results underscore the importance of hydrophobic matching for proper protein folding in biological systems and highlight how such physical features may be leveraged to enhance synthetic membrane-based materials. Specifically, leveraging physiochemical interactions of lipids and proteins between distinct membranes and within the hydrophobic region of single membranes will enable the design of membrane-based materials with enhanced transmembrane signaling (*9, 10, 14*) and more effective and specific engagement with cells (*13*), leading to more effective therapeutics.

This work is a major step towards utilizing *de novo* protein design, MD simulation, and membrane augmented cell-free systems to characterize complex biophysical phenomena, from the bottom up. Here, we show that protein hydrophobic mismatch alone can be used to enrich specific membranes with protein content and modulate protein-protein interactions. However, leveraging synthetic membranes and *de novo* designed proteins should enable the characterization of a variety of biophysical properties in a systematic, and controlled manner, and explore space not observed in natural systems. Finally, coupling *de novo* protein design with molecular dynamics highlights a powerful workflow which may enable a better understanding of protein biophysics and inform future protein design.

## Acknowledgments

We thank B. Liauw and A. Hunt for discussions on probing protein-protein interactions, and K. Warfel for discussing methods to assess protein expression. This research was supported in part through the computational resources and staff contributions provided for the Quest high performance computing facility at Northwestern University which is jointly supported by the Office of the Provost, the Office for Research, and Northwestern University Information Technology.

## Author Contributions

J. A. P., D. B., and N. P. K. conceived of the idea; J. A. P. and N. P. K. wrote the manuscript; P. L. and D. B. designed proteins; J. S. performed MD simulations, J. A. P., T. Q. V., T. F. G. designed and performed cell-free protein characterization experiments; all authors analyzed data, discussed results, and commented on the manuscript.

## Data and Materials Availability

All data can be found in the manuscript and supplementary files.

## Supplementary Materials

### Materials

1,2-dioleoyl-sn-glycero-3-phosphocholine (DOPC), 1,2-dimyristoleoyl-sn-glycero-3-phosphocholine (14:1 PC), 1,2-dierucoyl-sn-glycero-3-phosphocholine (22:1 PC), Cholesterol, 1,2-dipalmitoyl-sn-glycero-3-phosphocholine (DPPC), 1,2-dioleoyl-sn-glycero-3-phosphoethanolamine-N-(7-nitro-2-1,3-benzoxadiazol-4-yl)1,2-dioleoyl-snglycero-3-phosphoethanolamine-N-(lissamine rhodamine B sulfonyl) (18:1 Rhodamine), and 1,2-dioleoyl-sn-glycero-3-phosphoethanolamine-N-(Cyanine 5.5) (Cy 5.5 PE) were purchased from Avanti Polar Lipids. PURExpress and SNAP Alexa Fluor 488 were obtained from New England Biosciences. gBlocks and primers were ordered from Integrated DNA technologies and DNA was amplified and assembled using enzymes from Thermo Fisher. Phosphate-buffered Saline (PBS), sucrose, and Sepharose 4B (45−165 mm bead diameter) were obtained from Sigma Aldrich. Protein A/G beads, Calcein dye, streptavidin, Alexa Fluor 488 Biocytin, and rapamycin were purchased from Thermo Fisher. NanoBit substrate was purchased from Promega.

### Methods

#### Protein design

We designed transmembrane proteins with different transmembrane spans (with a range of 20-50 Å) by resurfacing the outside of the d*e novo* designer transmembrane proteins with patterned hydrophobic residues and adding RK- and YW-rings at the intracellular and extracellular boundary region, respectively. Briefly, hydrophobic residues are designed based on amino acid propensity in the membrane, replacing all polar residues exposed to the membrane. The design models of TMHC2 and TMH4C4 were used as the starting model.

**Table S1.**
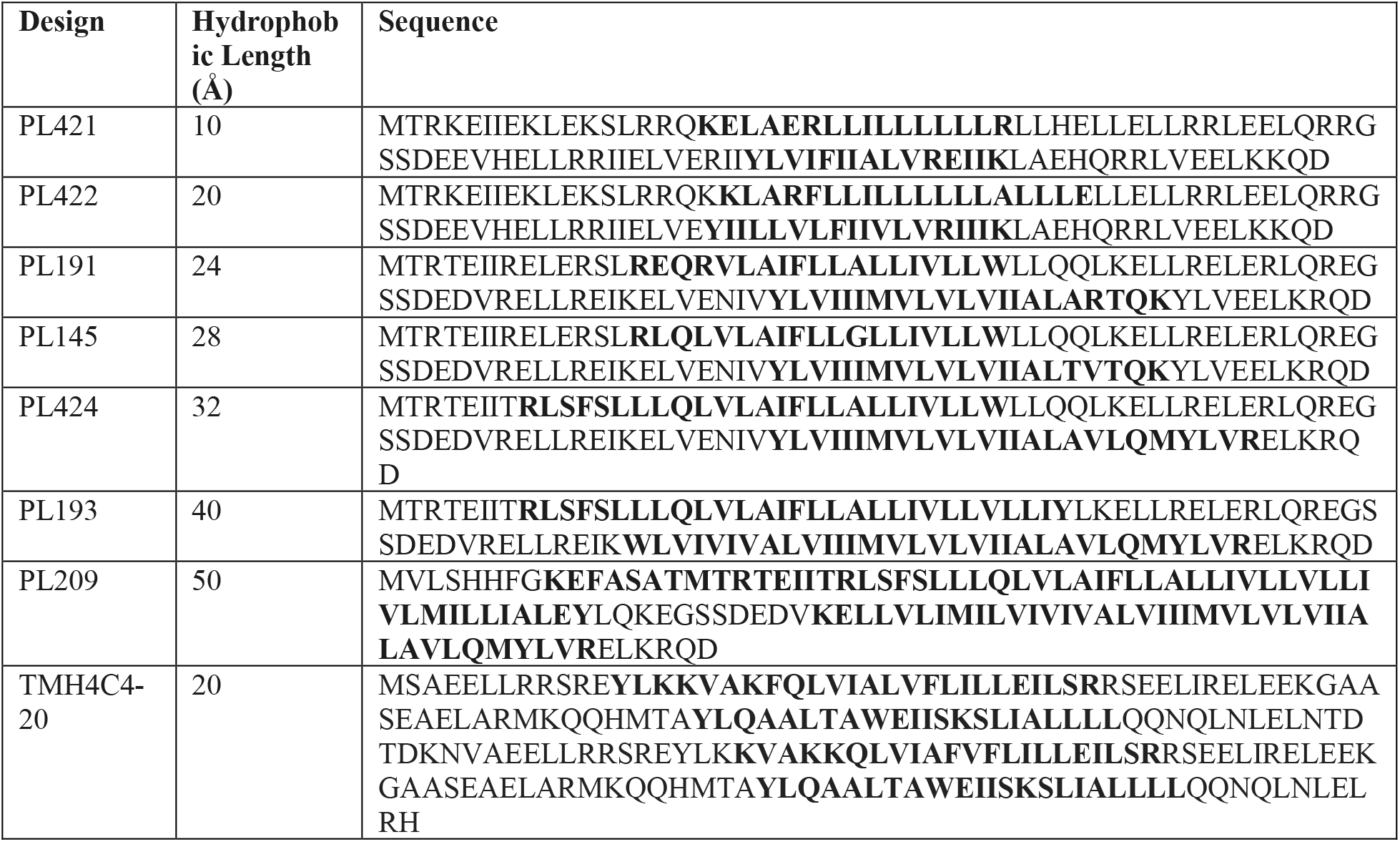
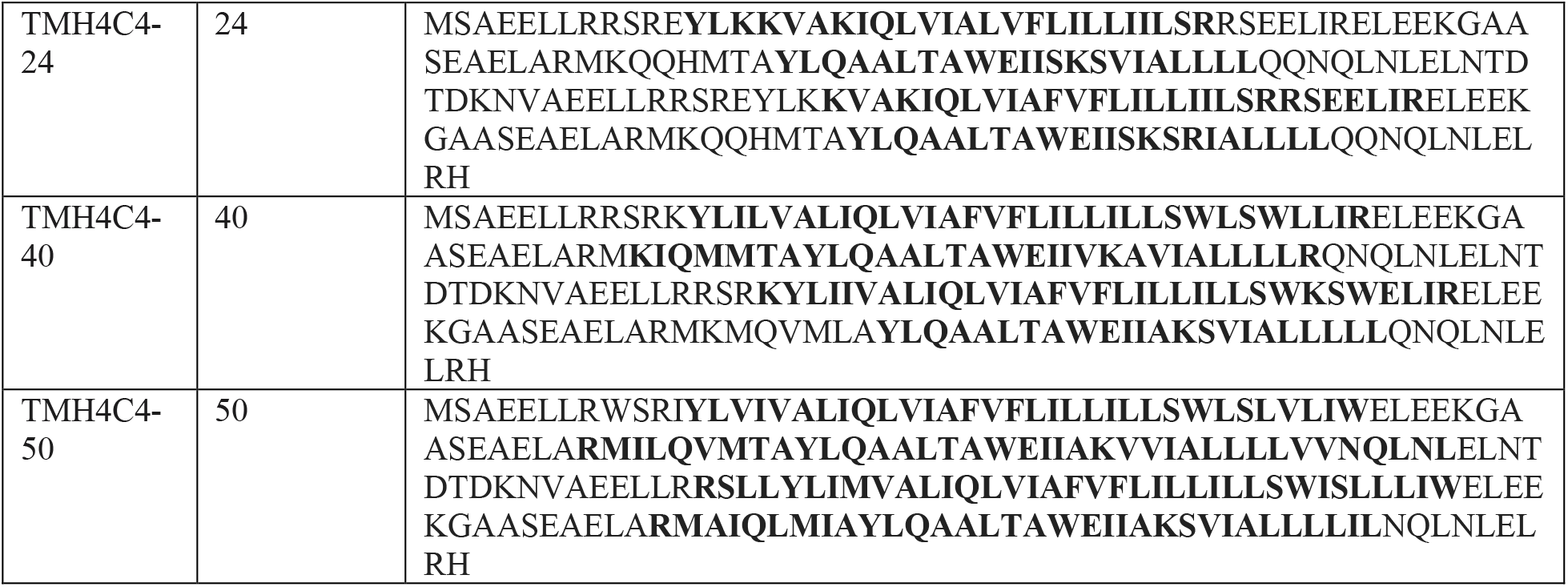
Sequences of protein designs. Transmembrane domains are marked in boldface.

#### Coarse grained simulations

Coarse-grained molecular dynamics simulations were conducted using the MARTINI force-field (v2.2) using GROMACS (2020.1). Simulations were performed using semi-isotropic pressure coupling to yield laterally tensionless membranes using the “martini straight” parameters (*34*).

The secondary structure of simulated protein was fixed in the simulations by an elastic network parametrized from the predicted protein structure (*35, 36*). Membranes of varying lipid compositions and protein assemblies were assembled using insane.py (*37*) and initially equilibrated for a minimum of 10 ns, productions runs were 6 μs with three replicates and sampled every 1 ns. If not indicated otherwise, the simulation was conducted at 295 °K. Trajectories were analyzed using MDAnalysis version 0.20.1 (*38*). Type of analysis and simulation size varying between systems: Data in panel Fig 1B, C was obtained for single component membranes with 216 MARTINI DYPC, DOPC or DGPC lipids per leaflet. The position in normal direction to the membrane of the PO4 bead (representing the phosphate headgroup) was analyzed around a single centered protein construct for both membrane leaflets. Then PO4 positions were binned by the radial distance from the protein center with 1 nm bin width. The difference between the PO4 position for each leaflet bin then determined the membrane thickness shown as an average over the whole trajectory. For simulations shown in Fig. 4A, membranes were composed of 138 DPPC, 92 DYPC and 99 cholesterol molecules per leaflet. In a radial selection around the protein center of mass, corresponding to the first layer of surrounding lipid molecules, individual lipid types were determined. The time average of detected lipids, then determined average membrane composition around the protein center at varying temperatures. For Fig. 3F the same DPPC:DYPC:cholesterol membranes as above were compared to membranes with 326 DOPC lipids per leaflet. Both membranes contained two copies of two different protein constructs. The distributions of center of mass protein-protein distances were determined for the two membrane compositions.

#### Gene Assembly and Cloning

Genes listed in SI Table S2 were ordered as gene blocks and cloned into a high copy plasmid used in previous work (*39*). Different fusion proteins were generated using standard restriction enzyme cloning techniques using Phusion DNA polymerase and restriction enzymes from Thermo Fisher. Pore proteins were toxic and prone to mutation and thus were not cloned into plasmids. Protein pores were ordered as gene blocks with elements required for gene transcription and translation (T7 promoter and terminator, ribosome binding site) and were amplified via PCR.

#### Vesicle Preparation

Throughout this study, vesicles were prepared via the thin film hydration method. Briefly, lipid was deposited into a glass vial and dried with a stream of nitrogen and placed under vacuum for 3 hours. Films were then rehydrated in Milli-Q water and heated at 60 °C for a minimum of 3 hours, and up to overnight. Vesicles were then briefly vortexed and extruded 21x through a 100 nm polycarbonate filter.

#### Analysis of folding and insertion of cell-free expressed proteins into synthetic membranes

Protein expression was performed with the PURExpress In Vitro Protein Synthesis kit (E6800, NEB) according to the manufacturer’s instructions. 30 μL reactions were assembled with a final concentration of 10 mM of lipid and 3.3 nM plasmid. Reactions were allowed to progress at 37 °C for 3 hours. GFP folding and fluorescence was monitored using a Molecular Devices Spectra Max i3 plate reader (ex 480 nm, em. 507 nm). Increase in GFP fluorescence was then calculated by subtracting the fluorescence at t=0 from the fluorescence at t=3 hours.

Protein expression was measured via western blot. Cell-free protein synthesis reactions were spun at 20,000 g for 10 minutes to pellet and remove uninserted protein. The supernatant was collected and run on a 12% Mini-PROTEAN TGX Precast Protein Gel (Bio-Rad) for all experiments, except the truncation experiments. For truncation experiments, samples were run on a 16.5% Tricine Mini-PROTEAN Precast Protein Gel to enhance the separation of smaller protein products. Wet transfer was performed onto a PVDF membrane (Bio-Rad) for 45 min at 100 V. Membranes were then blocked for an hour at room temperature in 5% milk in TBST (pH 7.6: 50 mM Tris, 150 mM NaCl, HCl to pH 7.6, 0.1% Tween) and incubated for 1 hour at room temperature or overnight at 4 °C with primary solution (anti GFP, diluted 1:1000 in 5% milk in TBST). Primary antibody solution was decanted, and the membrane was washed three times for 5 minutes in TBST and then incubated in secondary solution at room temperature for 1 hour (HRP-anti-Mouse (CST 7076) diluted 1:3000 in 5% milk in TBST). Membranes were then washed in TBST and incubated with Clarity Western ECL Substrate (Bio-Rad) for 5 min. Membranes were then imaged in an Azure Biosystems c280 imager and bands were quantified with ImageJ.

#### Preparation of Giant Unilamellar Vesicles

Giant, micron sized, vesicles were prepared via electroformation using the Nanion Vesicle Prep Pro (Nanion Technologies) standard vesicle preparation protocol. To visualize protein, proteins were expressed into liposomes containing 0.1 mol% 18:1 PC Cy5.5. Following expression, liposomes were diluted to 1 mM and 10 μL were deposited onto indium tin oxide slides and allowed to dry under vacuum for 30 minutes. Samples were then rehydrated with 290 mOsm sucrose. To visualize domains, 10 mM mixtures of lipid in chloroform were prepared with 0.1 mol% Rhod-PE. 10 μL of each solution was then drop-casted onto indium tin oxide slides and placed under vacuum for 20 minutes to eliminate solvent and rehydrated with 290 mOsm sucrose. GUVs were observed under a Nikon confocal microscope. Glass bottomed Lab-Tek II microscope chambers (Thermo Fischer) were used to image GUVs. 200 μL of bovine serum albumin was placed into each chamber and allowed to sit for 30 min. Each well was then washed with 290 mOsm PBS and 1 mL of 1 mM of GUVs were added to 250 mL of PBS and allowed to settle in each chamber. A 20x objective was used to visualize vesicles. Images were analyzed using NIS software.

#### Calcein Leakage

Vesicles were rehydrated with 50 mM Calcein in 10 mM HEPES. Calcein vesicles were purified using a size exclusion column packed with Sepharose 4B immediately before experimentation. PURExpress reactions were then assembled and calcein leakage was read (ex. 480 nm/ em. 520 nm) on the plate reader (Molecular Devices Spectra Max i3) for 3 hours at 37 °C. 1% Triton-X was then added to achieve a maximum dequenching of calcein, which served as the fluorescence intensity for 100% mixing. Percent content mixing was calculated using the following equation:

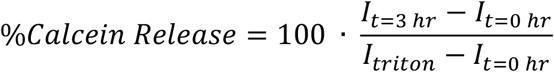

Where I_t=0hr_ is the initial fluorescence intensity, I_t=3hr_ is the fluorescence intensity at 3 hours, and I_triton_ is the fluorescence intensity after the addition of Triton-X. To determine the relative calcein release per protein, western blots were performed on samples. Calcein release values were then divided by total protein intensity for each sample to calculate the calcein release relative to protein expression.

#### Assessing protein sorting between distinct compartments via immunoprecipitation

100 nm 14:1 and 22:1 PC vesicles were prepared as outlined above with 0.1 mol% 18:1 PC Cy5.5 and 18:1 PC Rhodamine respectively. PURExpress reactions were assembled with 3.3 nM plasmid encoding either the 20, 24, 40, or 50 Å pore protein and 5 mM each of 14:1 and 22:1 PC vesicles. Reactions were allowed to progress at 37 °C for 3 hours. Samples were then incubated with Pacific-blue anti-flag antibody conjugated protein A/G beads for 1 hour at room temperature. Samples were washed 3 times and then analyzed via flow cytometry. Beads were gated for size (only larger beads were selected to eliminate unbound vesicles) and anti-flag antibody (405 nm excitation, 450/50 nm emission). Beads were analyzed for Rhodamine (550 nm excitation, 582/15 nm emission) and Cy5.5 (640 nm excitation, 730/45 nm emission with 685 longpass filter). At least 10,000 events were recorded, and beads were re-gated in FlowJo (TreeStar).

#### Analyzing differential pore activity

14:1 and 22:1 PC vesicles were prepared as outlined above with 0.1 mol% 18:1 PC Cy5.5 and 18:1 PC Rhodamine respectively. Lipid films were rehydrated with 5 μM streptavidin and extruded to 1 μm. PURExpress reactions were assembled with 3.3 nM plasmid encoding either the 20, 24, 40, or 50 Å pore protein and 5 mM each of 14:1 and 22:1 PC vesicles and incubated at 37 °C for 3 hours. Reactions were then purified via size exclusion chromatography to purify away unencapsulated streptavidin. Vesicles were incubated with 1 μM biocytin conjugated Alexa Fluor 488 for 24 hours. Samples were diluted to a lipid concentration of 1 μM in PBS and analyzed via flow cytometry on a BD LSR Fortessa Special Order Research Product (Robert H. Lurie Cancer Center Flow Cytometry Core). Alexa Fluor 488 was excited with a 488 nm laser and captured with a 505 nm long pass filter and a 530/30 nm bandpass filter, Rhodamine was excited with a 552 nm laser and captured with a 582/15 nm bandpass filter, and Cy5.5 was excited with a 640 nm laser and captured with a 685 nm longpass filter and a 730/45 nm bandpass filter. Events on the cytometer were thresholded on the presence of either Rhodamine or Cy5.5 detection to identify vesicles, and approximately 100,000 events were captured per reaction. Data was analyzed in FlowJo v10.8 and spectrally compensated. Samples were gated using curly quad gating of Rhodamine versus Cy5.5 to isolate single-dye positive events and thus restrict analysis to only thin or thick membrane vesicles (Fig. S9). Curly quad gating (rather than quad gating) was necessary to account for photon counting and measurement error at high laser settings ((*40*). Single lipid-dye positive events were then gated for Alexa Fluor 488 using samples containing vesicles but no Alexa Fluor 488. The percent of thin and thick vesicles identified as Alexa Fluor 488 positive and the mean fluorescent intensity (MFI) of each vesicle population was determined and analyzed. Vesicles that were prepared as above but that did not have a protein pore (i.e., the PURExpress reaction did not have pore-encoding DNA) was used to determine background. This background signal (e.g., percent positive vesicles or MFI) was subtracted as part of the data analysis process.

#### Lipid-Protein FRET experiments

Vesicles composed of DOPC or 42.5 mol% 14:1 PC/27.5 mol% DPPC/30 mol% cholesterol were prepared with 0.1 mol% 18:1 PC Rhodamine as outlined above. PURExpress reactions were prepared with 10 mM vesicles and 3.3 nM plasmid encoding the 20, 24, 40, or 50 Å hairpin proteins with a C-terminal SNAP tag. Reactions were performed at 37 °C for 3 hours. Samples were then incubated with 10 μM Alexa Fluor-SNAP substrate for 30 minutes at 37 °C. Vesicles were purified away from free SNAP substrate via size exclusion chromatography. Vesicles were collected and FRET was measured using an Agilent Cary Eclipse Fluorescence Spectrophotometer by exciting the samples at 488 nm and recording the emission at 520 and 590 nm. Fluorescence measurements were recorded at temperatures ranging from 25 to 47 °C. Vesicle samples were then treated with trypsin and 0.1% Triton X to disrupt vesicles and SNAP conjugated dye.

Relative FRET, noted here as C_D_/C_H_, was calculated using the following equation:

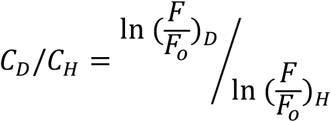

Where F is the fluorescent intensity of donor in the presence of acceptor, F_o_ is the fluorescent intensity of donor after the addition of trypsin and Triton-X. D denotes samples with domain forming membrane and H denotes samples with homogenous membranes. With this convention, C_D_/C_H_ will be high if Rhodamine (acceptor) and protein (donor) partition into the same lipid domain and low if they are segregated into different lipid domain (*41*).

#### NanoBit Experiments

Vesicles composed of DOPC or 42.5 mol% 14:1 PC/27.5 mol% DPPC/30 mol% Cholesterol were prepared as outlined above and extruded to 100 nm. PURExpress reactions were assembled with 1.7 nM of each DNA construct: 20 Å Hairpin/ 50 Å Hairpin, 20 Å Hairpin/ 20 Å Hairpin, 50 Å Hairpin/ 50 Å Hairpin. Reactions were allowed to progress for 3 hours at 37°C.

For rapamycin experiments, cell-free reactions were split into two and either rapamycin in DMSO or DMSO only was added to protein incorporated vesicles at a final concentration of 30 nM (or a DMSO mol fraction of 1 mol lipid: 0.0015 mol rapamycin). Samples were incubated for 2 hours at room temperature. NanoBiT reactions were setup using the Promega Nano-Glo Live Cell Assay System following the Technical Manual with minor modifications. Cell free reactions were diluted 1:4 in 1x PBS and the Nano-Glo Substrate was used at a 50x final dilution of the stock. Luminescence was read using a Molecular Devices Spectra Max i3 plate reader at room temperature for 10 minutes. To ensure the ratios of NanoBit to Substrate were in optimal range, luminescence was checked to be constant over the ten-minute read. Rapamycin induced luminescence was then calculated as:

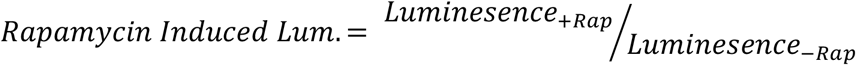

Where luminescence_+Rap_ is the measured luminescence in the presence of rapamycin and luminescence_-Rap_ is the measured luminescence in the presence of DMSO only.

To characterize protein-protein interactions with increasing temperature, the luminescence of samples was then recorded at varying temperatures from room temperature to 45 °C. Relative NanoBit assembly was then calculated as:

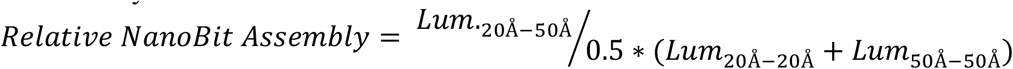

Where Lum_20Å-50Å_ is the luminescence of samples with 20 Å and 50 Å hairpin proteins, Lum._20Å-20Å_ is the luminescence of samples with 20 Å and 20 Å hairpin proteins, and Lum._50Å-50Å_ is the luminescence of samples with 50 Å and 50 Å hairpin proteins. Luminesce values were then normalized to the luminesce value at room temperature. Dividing by the average of NanoBit fused to proteins of the same length allows for the increase in Nanobit assembly due to increases in lipid and protein mixing as systems with the same TMDs should reside in the same lipid domains. Furthermore, this normalization accounts for luminescence differences due to temperature.

## Supplemental Figures

**Figure S1.**
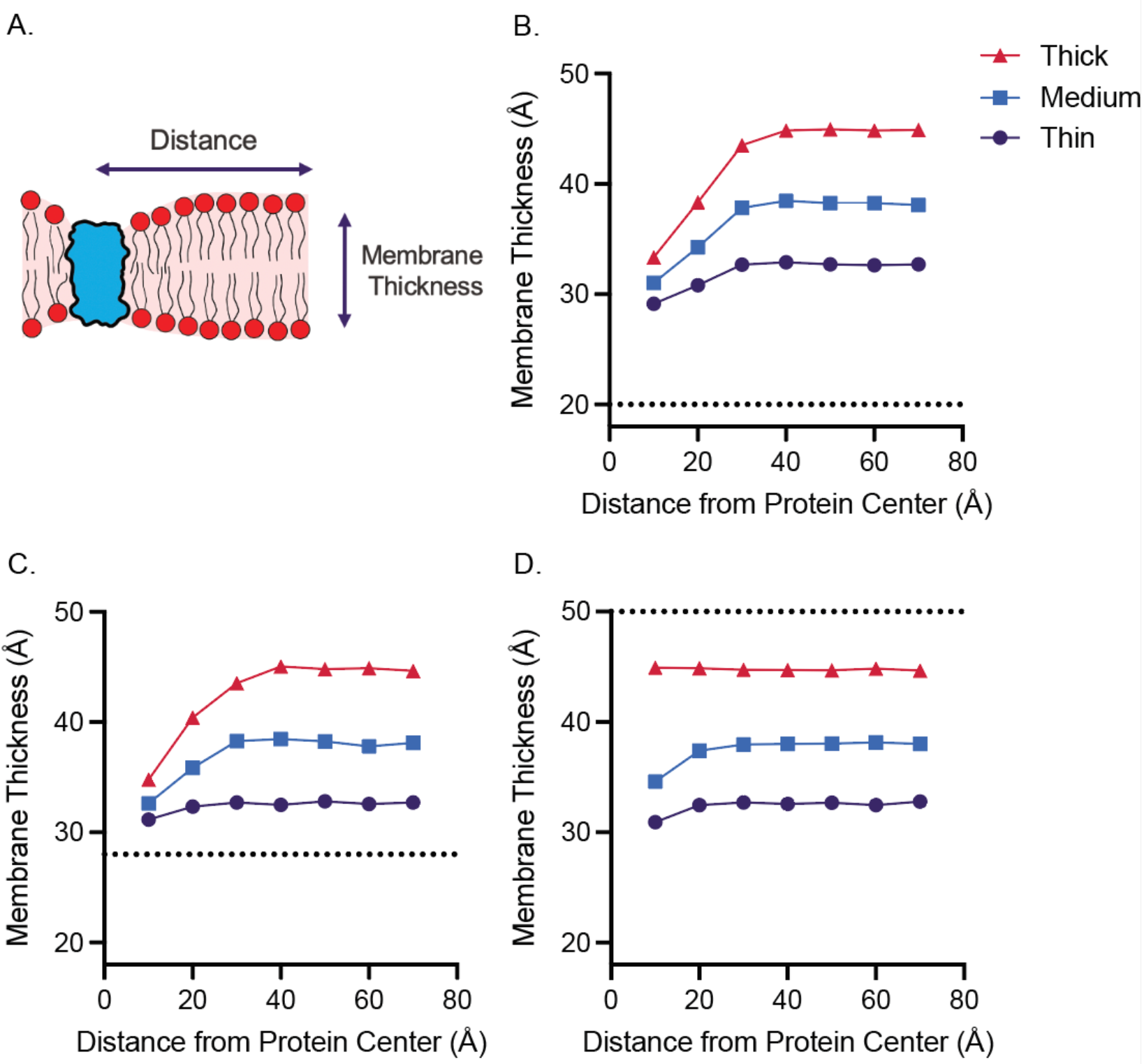
Membranes must deform to accommodate hydrophobic mismatch. Using MD simulations, the thickness of the membrane was recorded as a function of distance from the protein (**A**). This analysis was performed for the 20 (**B**), 28 (**C**), and 50 Å (**D**) proteins. Horizontal dotted lines represent the hydrophobic thickness of the protein. Data presented in Fig. 2C was generated by subtracting the membrane thickness at 70 Å from membrane thickness at 10 Å from the protein center.

**Figure S2.**
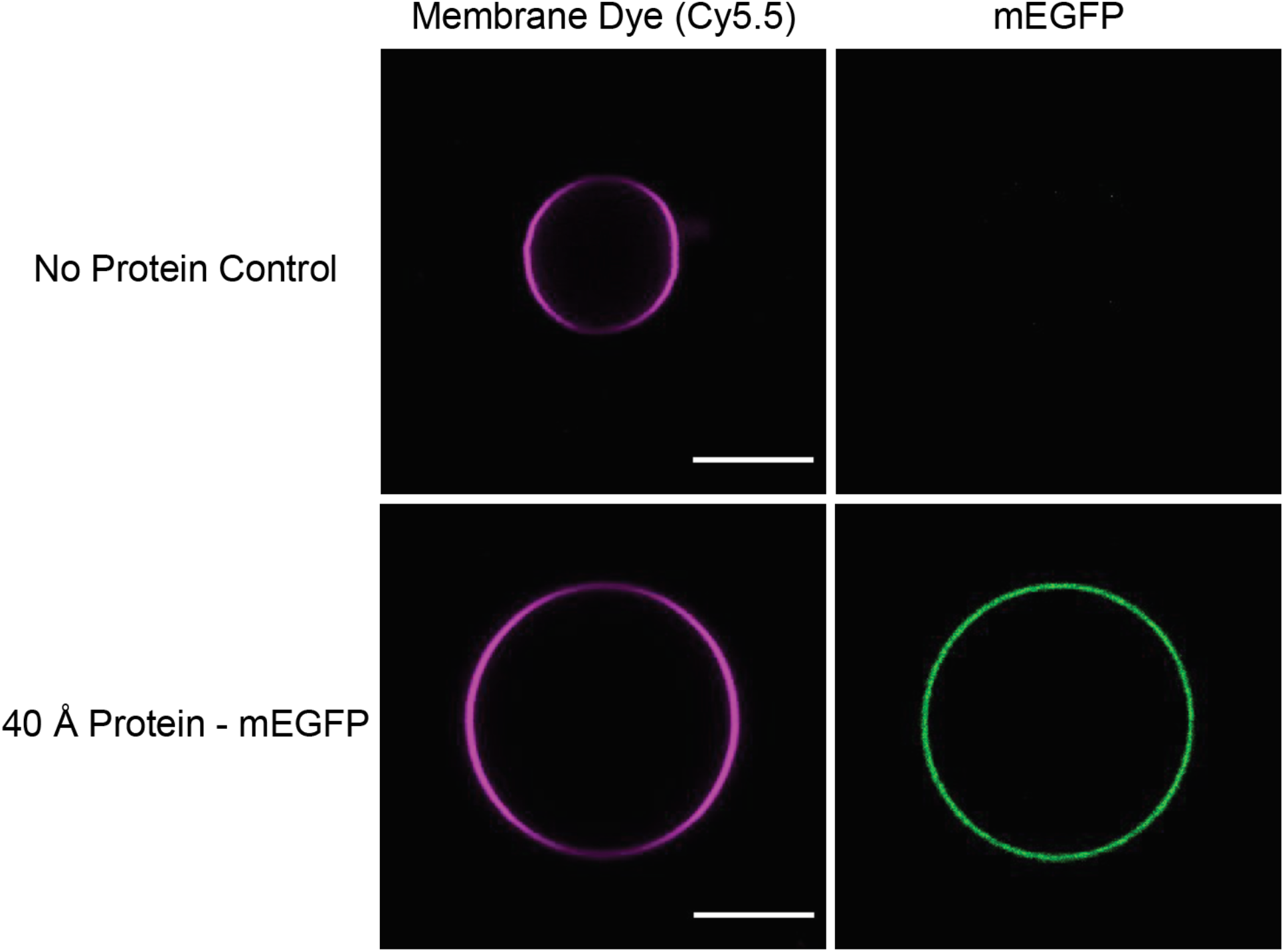
Membrane proteins insert in fold into synthetic membranes as assessed by confocal microscopy. GUVs were prepared with Cy5.5 conjugated lipid (left column). When proteins are expressed into synthetic lipids, GFP localizes to the membrane, indicating membrane protein insertion and folding (right). Pictured is the 40 Å protein in a DOPC membrane. Scale bar 10 μm.

**Figure S3.**
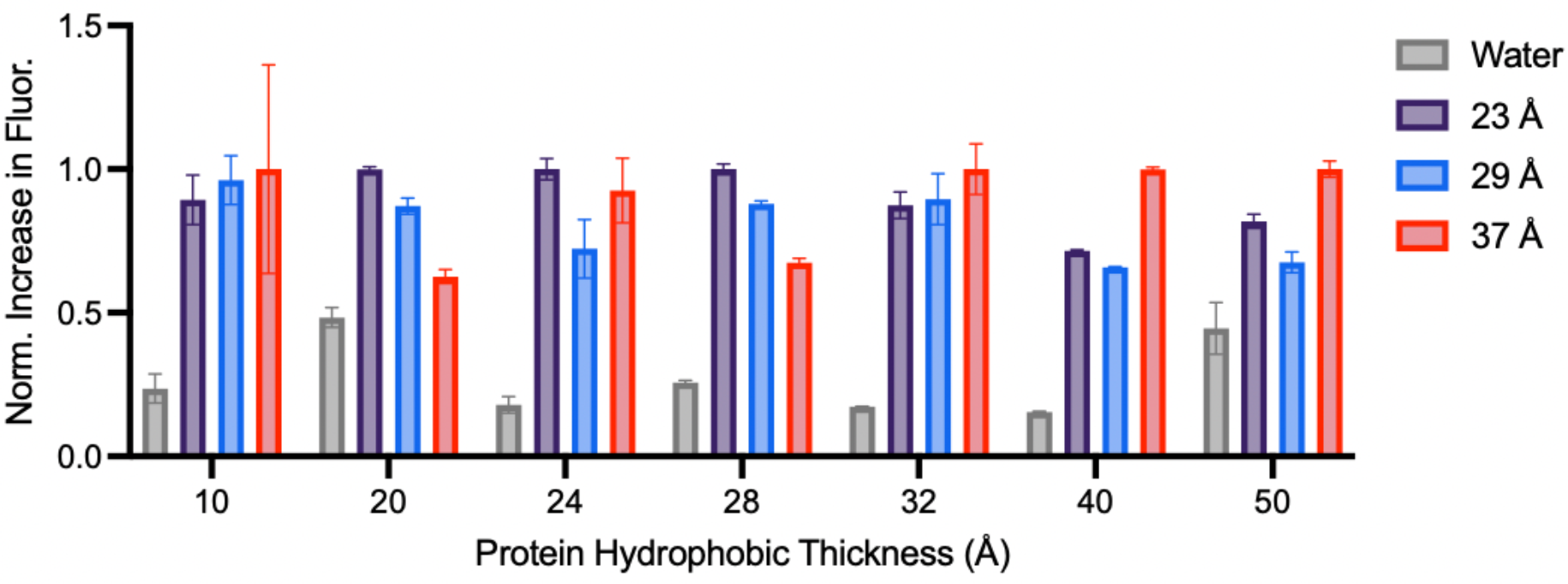
Highest increases in GFP fluorescence are observed when hydrophobic mismatch is minimized. Data presented in Figure 2D replotted as a bar graph to display error. Increase in GFP fluorescence was normalized to the maximum increase for each construct. *n=3*, error bars represent S. E. M.

**Figure S4.**
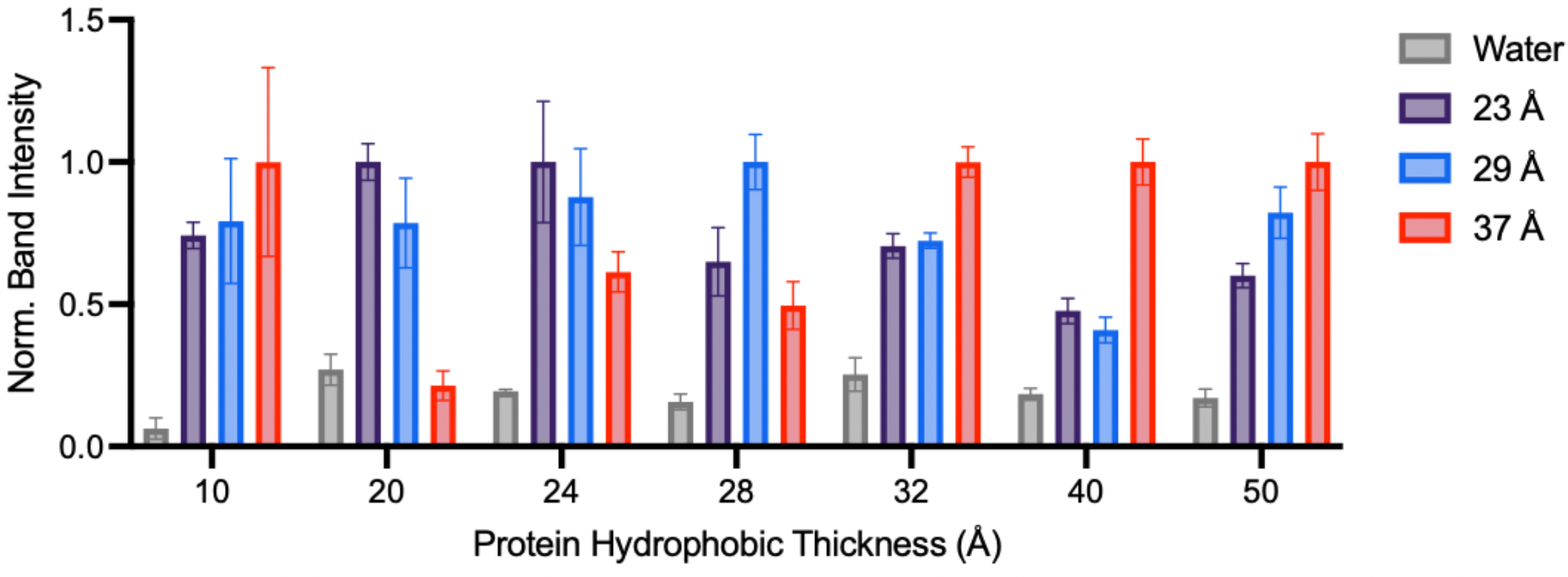
Highest protein expression levels are observed when hydrophobic mismatch is minimized. Western blots were performed on samples represented in Fig. 1D. Protein band intensity was normalized to the most intense band for each construct. *n=3*, error bars represent S. E. M.

**Figure S5.**
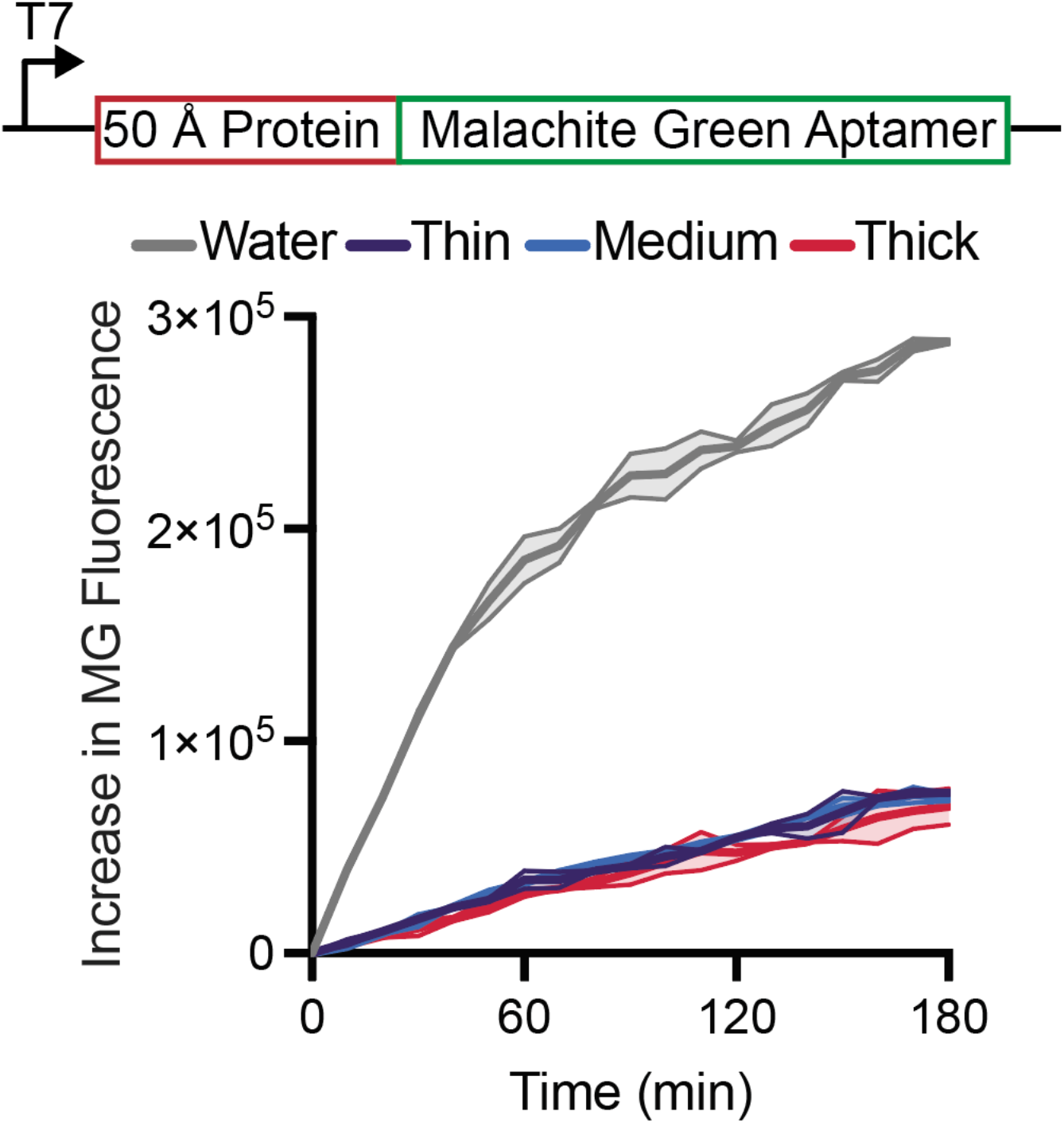
Transcription does not change as a function of membrane-protein hydrophobic mismatch. Transcription of the 50 Å protein, as reported by malachite green fluorescence, decreases with the addition of lipid vesicles, but does not change as a function of hydrophobic mismatch. *n=3*, error bars represent S. E. M.

**Figure S6.**
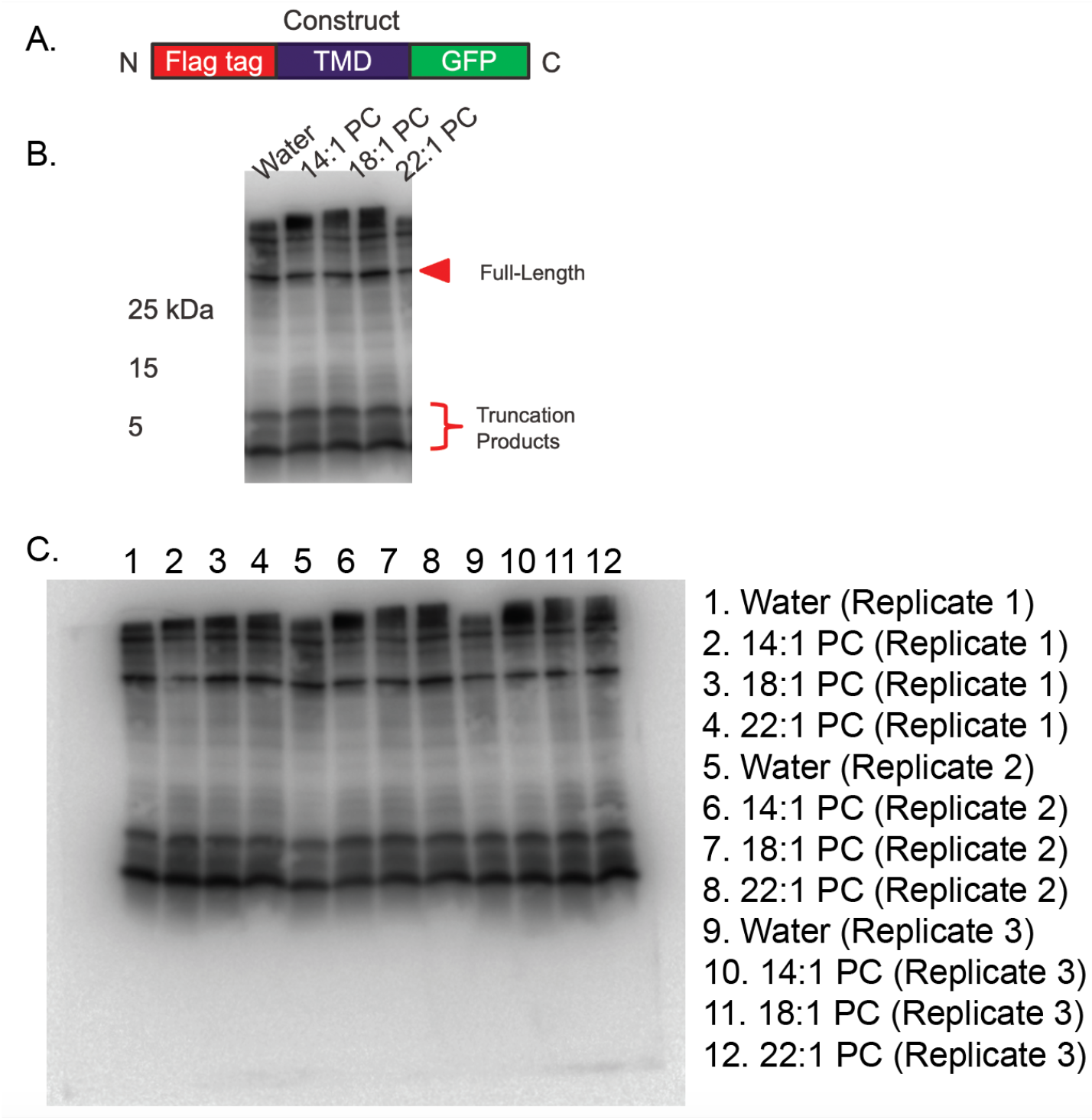
Analysis of truncation products via western blot. (**A**) An N-terminal flag tag was added to the 50 Å hairpin protein enabling the detection of all protein products. (**B**) The construct was expressed in the presence of water, 14:1 PC, 18:1 PC, and 22:1 PC and protein formation was assessed via western blot. The intensity of the full-length product and truncation products was measured using ImageJ and the ratio of full-length to truncation product intensity was plotted in Fig. 2F. Uncropped western blot for all replicates is displayed in (**C**).

**Figure S7.**
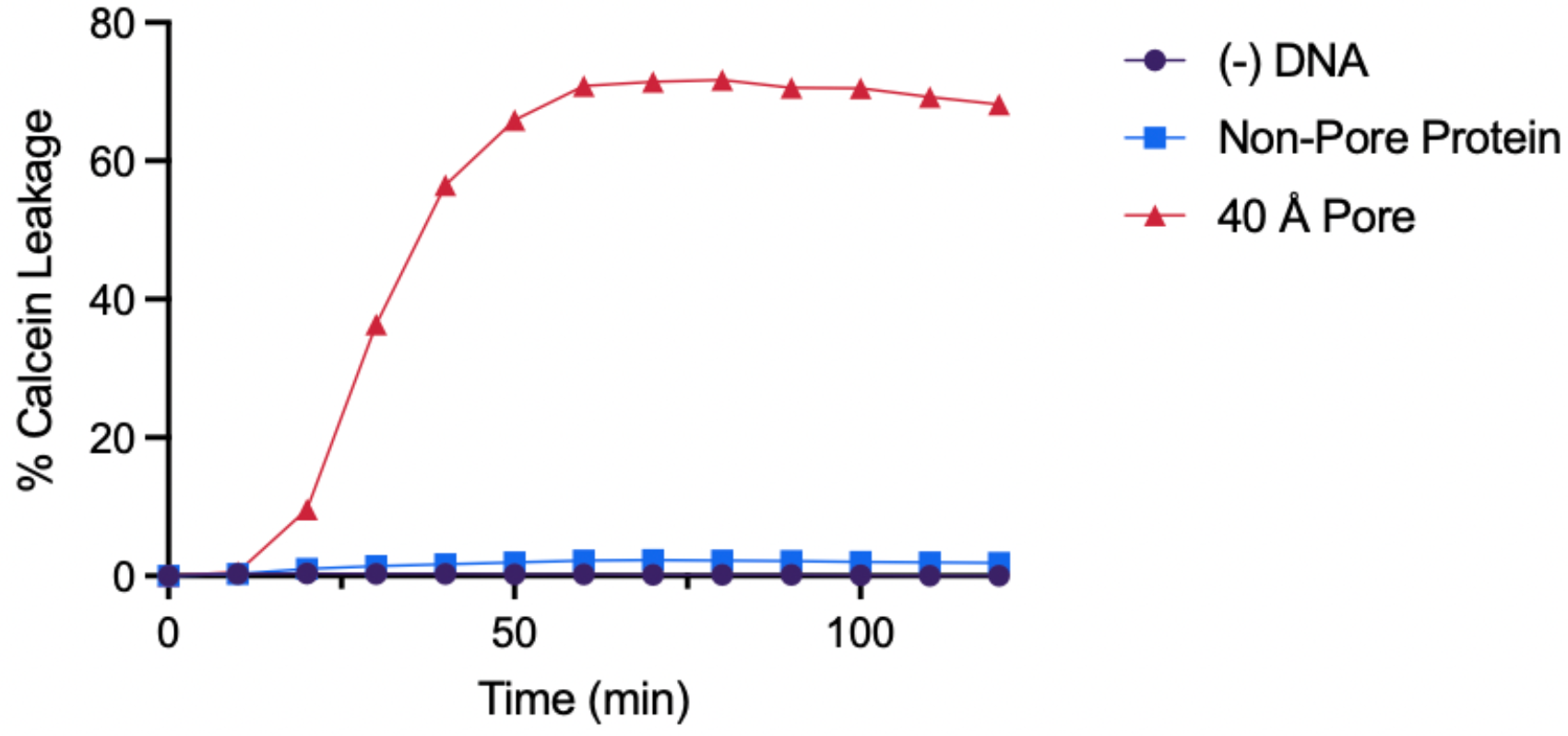
The expression and insertion of transmembrane pore proteins enables the release of calcein dye. 50 mM calcein, a self-quenching dye, was encapsulated into DOPC vesicles. Vesicles were then added to a cell free reaction without DNA, or DNA encoding the 40 Å hairpin or pore protein. Upon expression and insertion of the 40 Å pore protein, an increase in fluorescence because of calcein leakage and subsequent dequenching was observed. This demonstrates that calcein leakage is specific to pore protein expression and integration into vesicle membranes, and not due to interactions with PURExpress or insertion of non-pore proteins.

**Figure S8.**
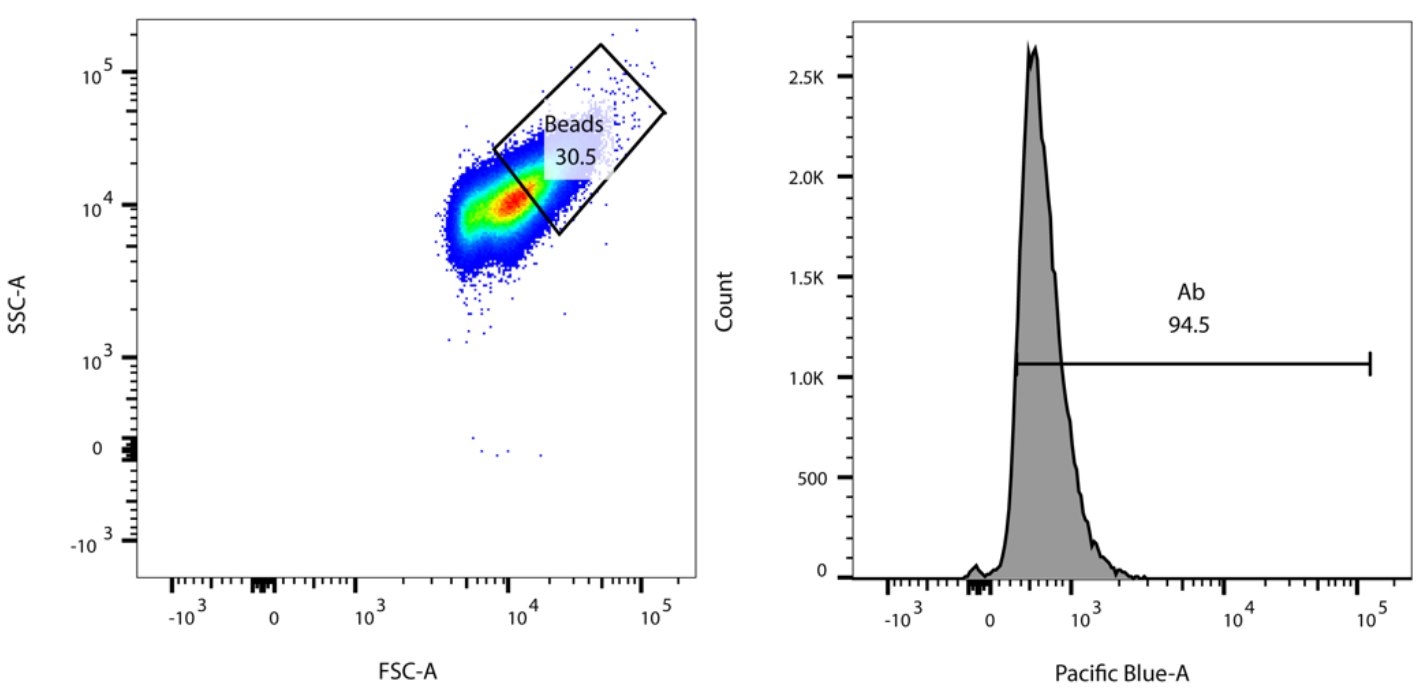
Gating strategy for bead-based protein sorting assays (Figure 3C and 3D). Beads were first gated on sizes (left) to analyze larger beads and reduce analysis of unbound vesicles, then gated by the presence of antibody (right).

**Figure S9.**
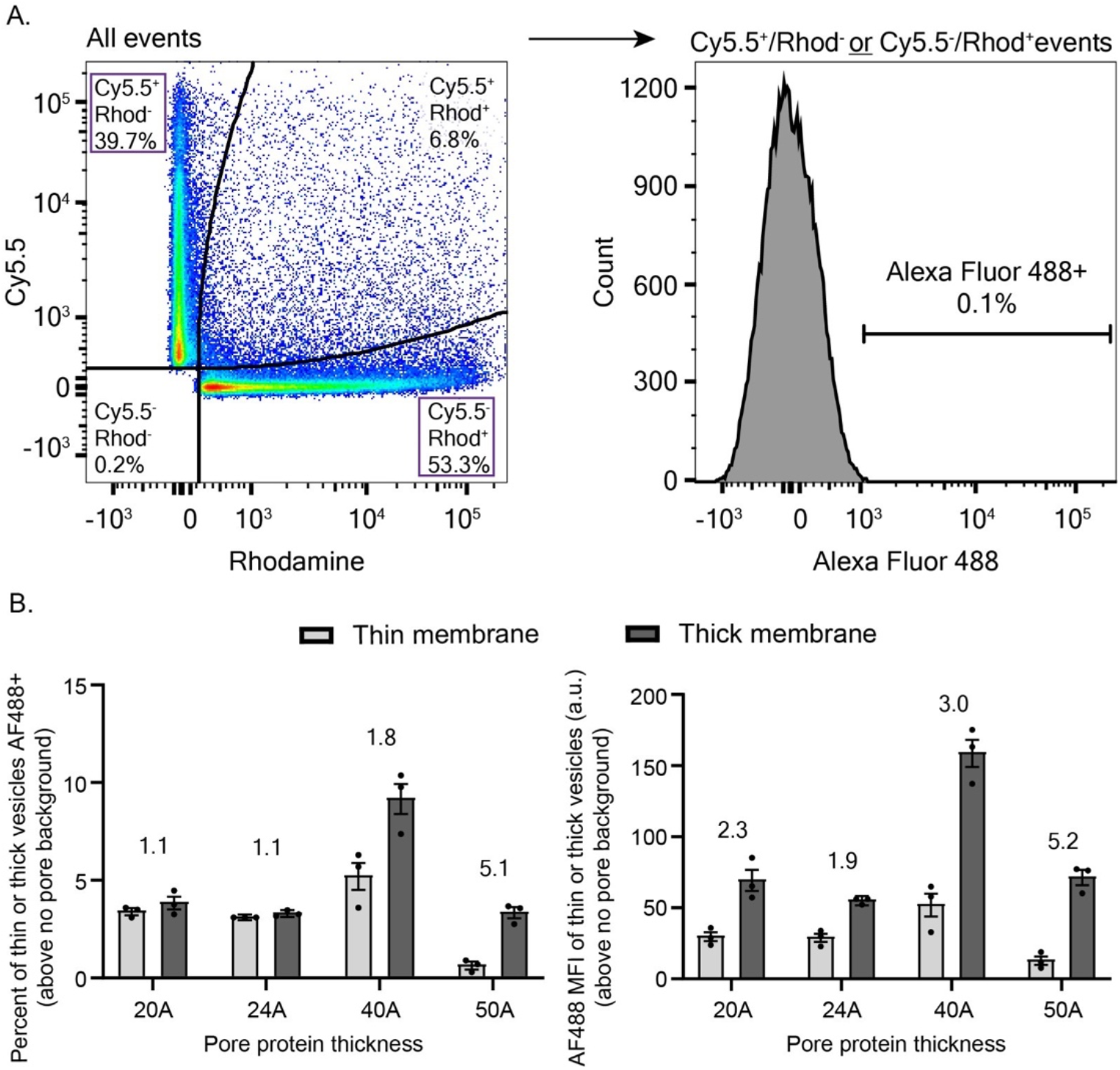
Gating strategy and population metrics for pore permeability flow data presented in Figure 3E and 3F. Thin and thick membrane vesicles were labeled with a single membrane dye, either Cy5.5 or Rhodamine, respectively. (**A**) To analyze subsequent flow cytometry data, events were first gated via a curly quad gating strategy. This identified vesicle populations with only a single membrane dye (left image, top left gate and bottom right gate). A negative control sample without Alexa Fluor 488 dye was used to set the threshold for an Alexa Fluor 488 (AF488) positive signal (right). This gate was then applied to single positive vesicle populations to determine Alexa Fluor 488AF488 signal as a measure of functional protein pore insertion. (**B**) Population level metrics of the data presented in Figure 3F. The left figure depicts the percentage of vesicles with a single membrane dye that are also AF488 positive per the gating strategy in (**A**) for different protein pore thicknesses. The right figure depicts the AF488 mean fluorescence intensity (MFI) of vesicles with a single membrane dye for a given pore protein thickness (i.e., the MFI is calculated from samples gated per only the left image in (**A**). In both cases, the data are background subtracted; the background was determined from vesicles incubated with biocytin but not a co-expressed pore protein. Data points represent the three replicates for each condition, the bar graphs represent the mean, and the error bars represent the standard deviation. The numbers are ratios of thick membrane average to thin membrane average for a given metric and pore protein thickness. We take these ratios to

**Figure S10.**
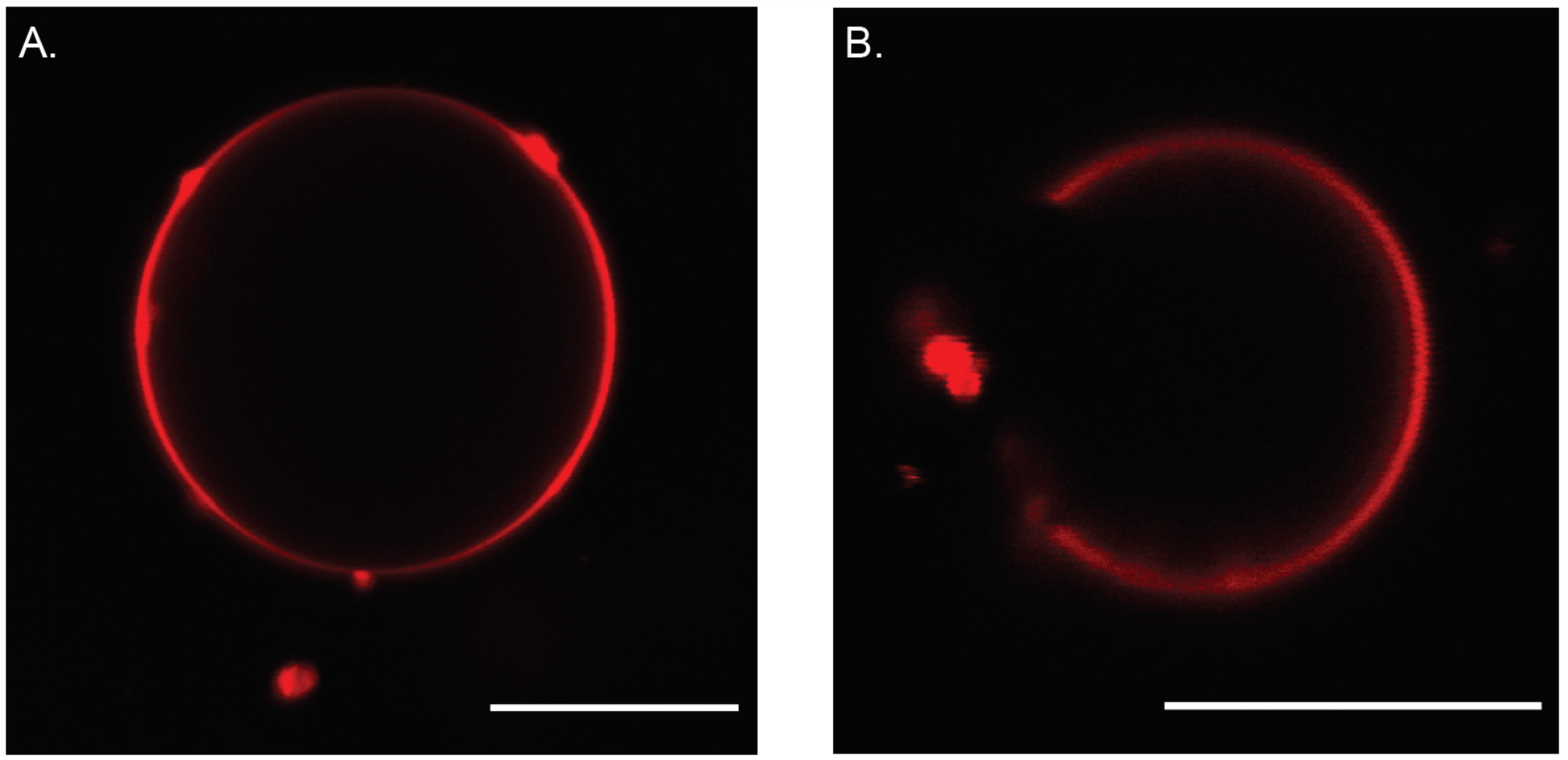
Fluorescent microscopy of giant unilamellar vesicles demonstrates that lipid mixtures do not form microdomains. (**A**) Vesicles composed of 42.5 mol% 14:1 PC/27.5 mol% DPPC/30 mol% Chol (composition used in this study) and (**B**) 40 mol% 14:1 PC/40 mol% DPPC/20 mol% Chol. Membranes were labeled with 0.1 mol% 18:1 PC Rhodamine, which localized to the lipid disordered phase. Absence of dye, as seen in (**B**), indicates the presence of microdomain formation, a property often selected in previous studies. By increasing the cholesterol content and decreasing the amount of DPPC, membranes that do not exhibit microdomain formation are formed. Samples with higher cholesterol content do not exhibit microdomain formation. Scale bars are 10 μm.

**Figure S11.**
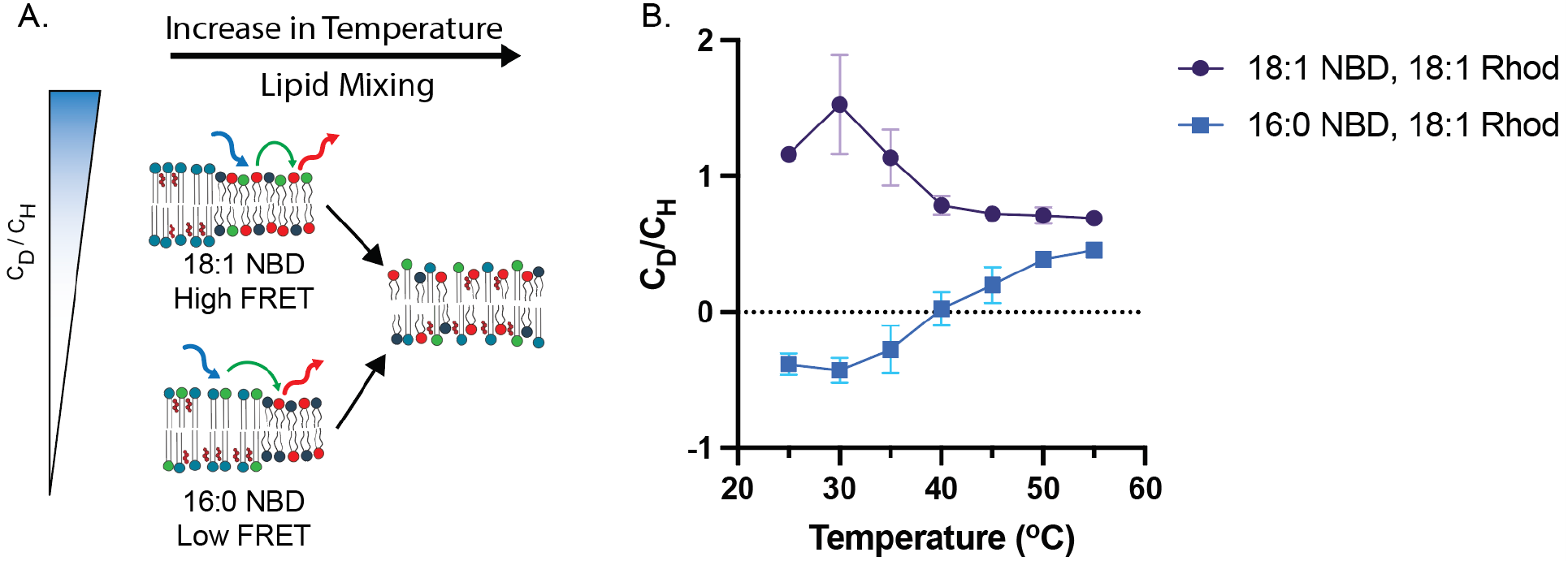
Lipid organization can be detected by lipid-lipid FRET. (**A**) Differences in C_D_/C_H_ can be measured when incorporating either 18:1 or 16:0 conjugated NBD (ex. 460 nm /em. 535 nm) in vesicles composed of 42.5 mol% 14:1 PC/27.5 mol% DPPC/30 mol% Cholesterol and 18:1 PC Rhodamine (ex. 560 nm/em. 590 nm). Lipid-lipid FRET was used to validate C_D_/C_H_ as lipid phase separation has been well characterized and is more well defined. (**B**) 18:1 PC NBD resides in the liquid disordered lipid phase, with 18:1 Rhodamine, and this has a higher C_D_/C_H_ compared to vesicles with 16:0 NBD, which resides in the liquid ordered phase, farther from 18:1 Rhodamine (lower C_D_/C_H_). Upon increasing temperature, C_D_/C_H_ converge, indicated that lipids intermix at higher temperatures due to increases in fluidity and subsequent lipid miscibility.

**Figure S12.**
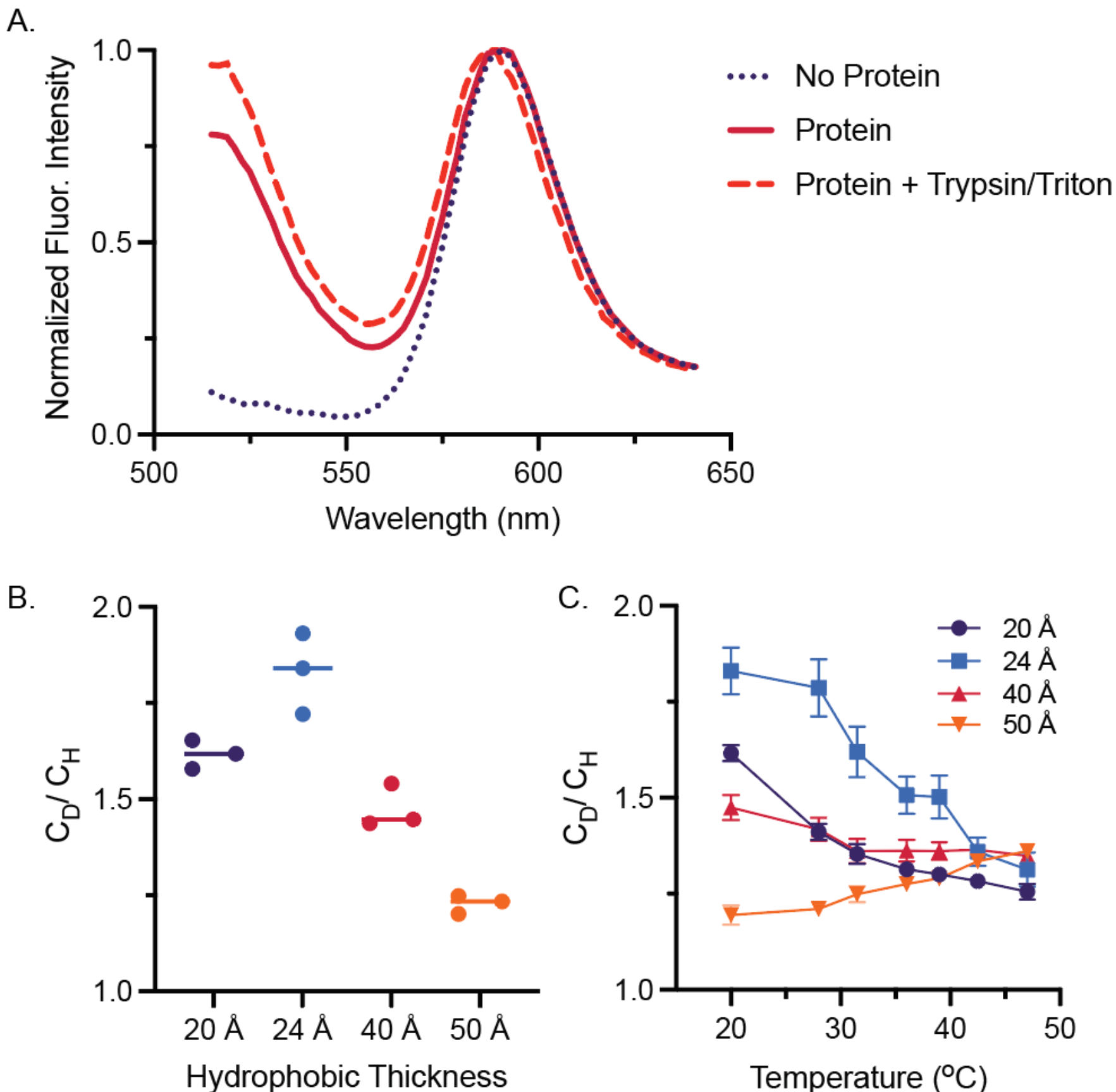
SNAP conjugated fluorophores enable lipid-protein FRET and protein-lipid interactions to be probed in vitro. (**A**) Spectra of SNAP conjugated protein in a homogenous DOPC membrane containing 0.1 mol% 18:1 Rhodamine before and after the addition of trypsin and triton. The dequenching of AF488 after the addition of triton and trypsin indicates that lipid-protein FRET is occurring. (**B**) At room temperature, C_D_/C_H_ values are higher for membrane proteins with shorter transmembrane domains, indicating that the shorter the transmembrane domain, the closer the protein is to rhodamine on average. (**C**) Upon increasing temperature, C_D_/C_H_ for all constructs converges to the same value, indicating that lipid and proteins mixing can occur at elevated temperatures.

**Fig S13.**
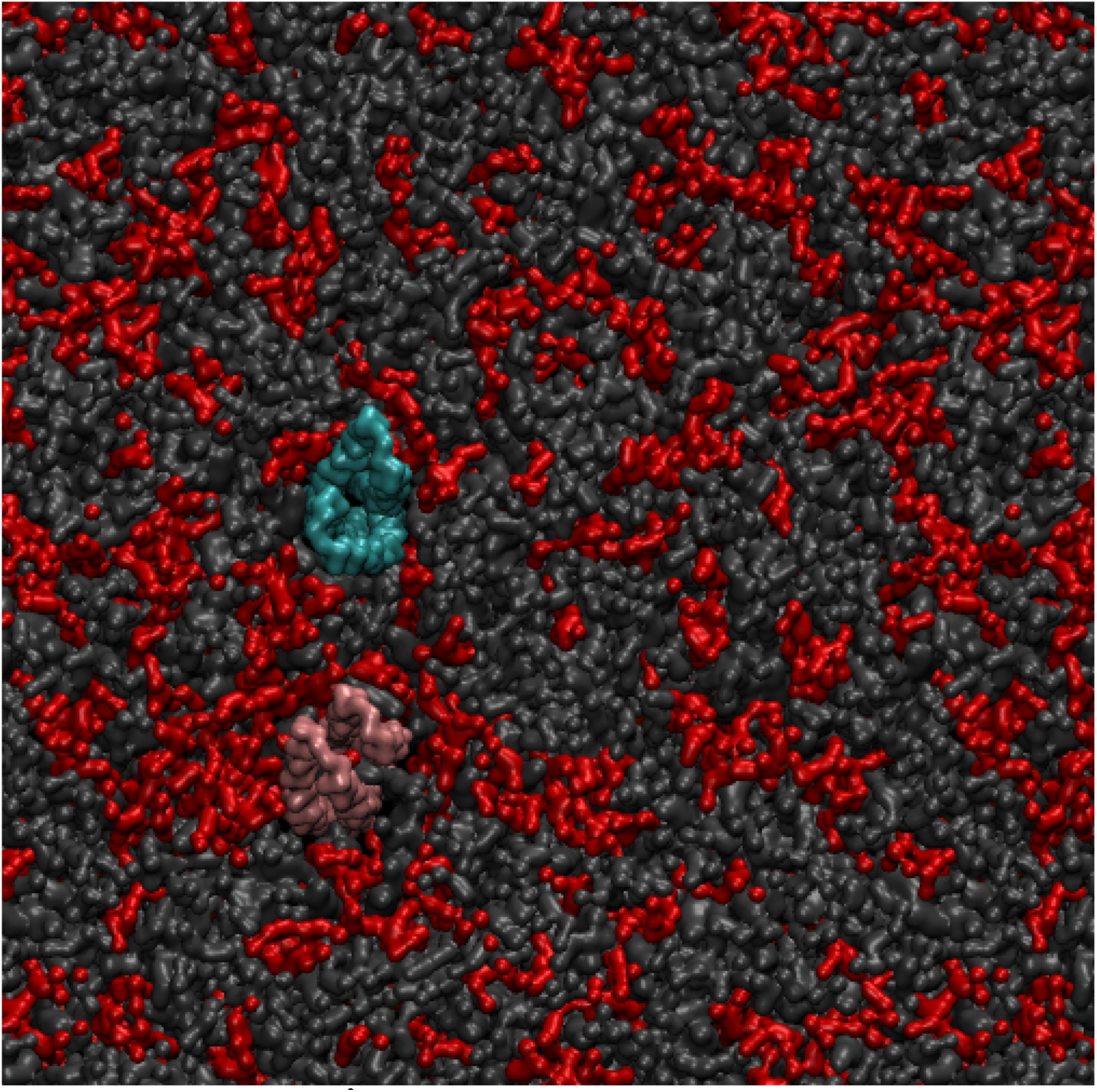
Simulation of 20 and 50 Å proteins in phase separating membranes. DYPC lipid (red) nucleates around the 20 Å hairpin protein (pink). The 50 Å protein (blue) is in contact with DPPC and cholesterol (grey) more often than DYPC. Further, proteins are apart from one another. Representative image is from Movie S1. The membrane is composed of 42.5 mol% 14:1 PC/27.5 mol% DPPC/30 mol% Cholesterol.

**Figure S14.**
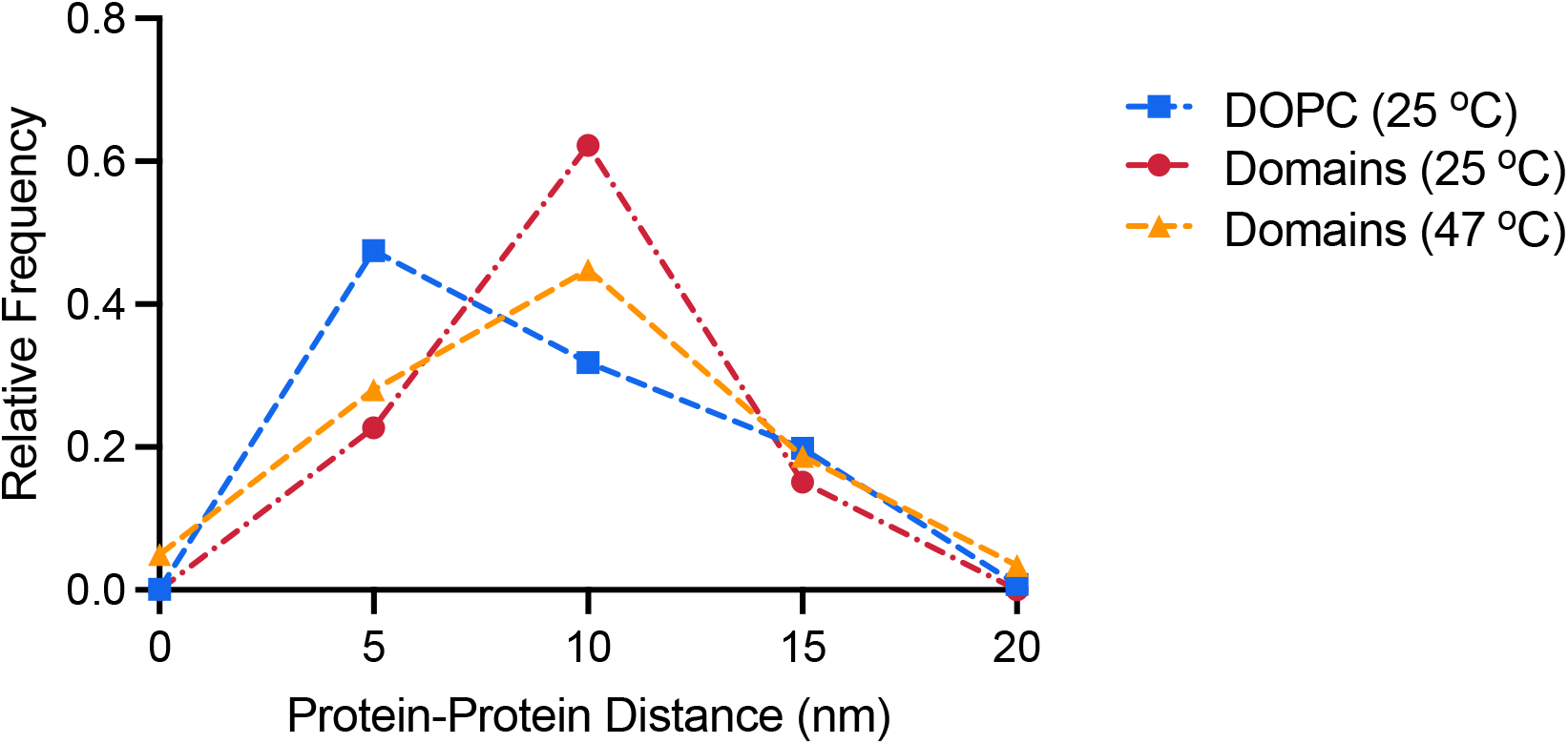
MD simulations predict decreased protein-protein distance at elevated temperatures. At room temperature, protein-protein distance between the 20 and 50 Å hairpin is on average smaller in homogenous, single component DOPC membranes (**A**) compared to membranes composed of 42.5 mol% 14:1 PC/27.5 mol% DPPC/30 mol% Cholesterol (**B**). (**C**) At 47°C, protein-protein distance decreases in membranes composed of 42.5 mol% 14:1 PC/27.5 mol% DPPC/30 mol% Cholesterol due to increased lipid mixing. Histograms were generated from 3 independent simulations.

**Figure S15.**
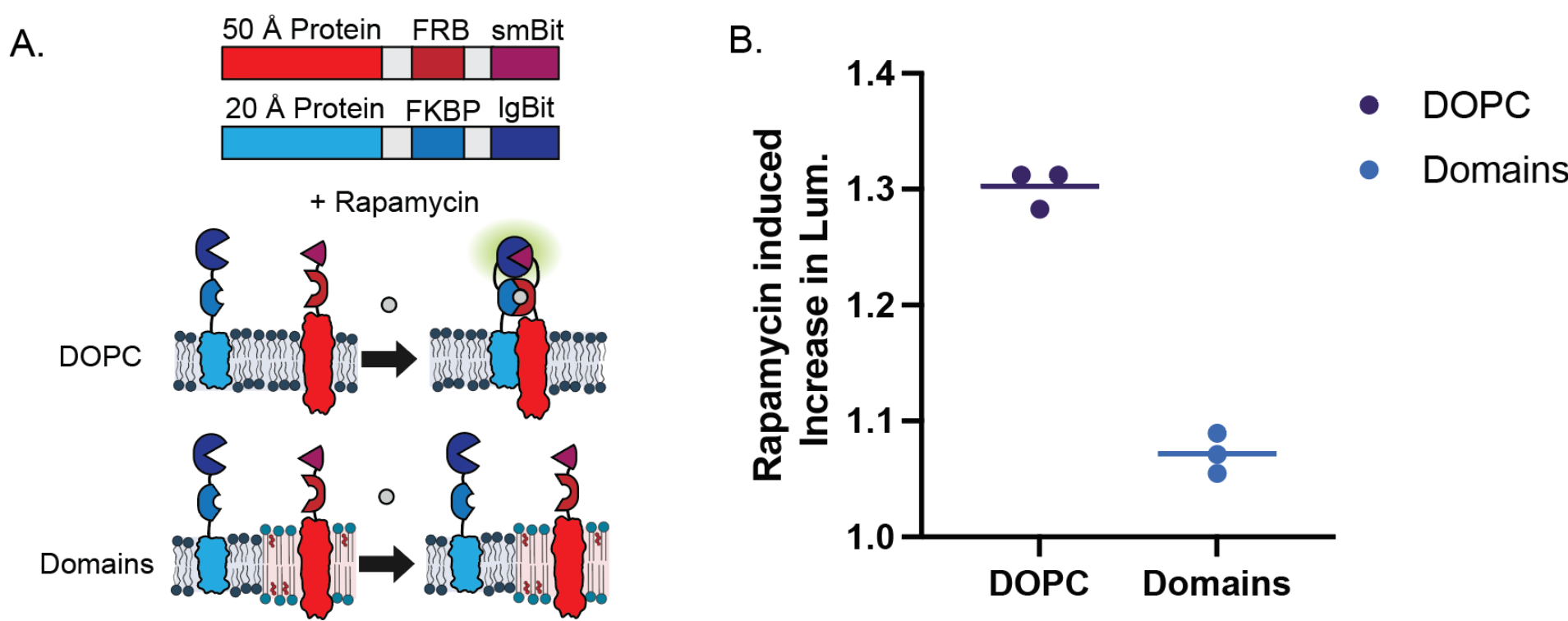
Proteins are more prone to dimerization in DOPC membranes compared to membranes prone to phase separation. (**A**) FRB and FKBP were fused to the C-terminus of the 20 and 50 Å protein, respectively. Addition of rapamycin forced proteins to dimerize and NanoBit to become reconstituted, enabling the evaluation of protein-protein interactions. (**B**) An increase in luminescence, because of the protein dimerization, was observed in DOPC membranes, but not membranes prone to domain formation.

**Figure S16.**
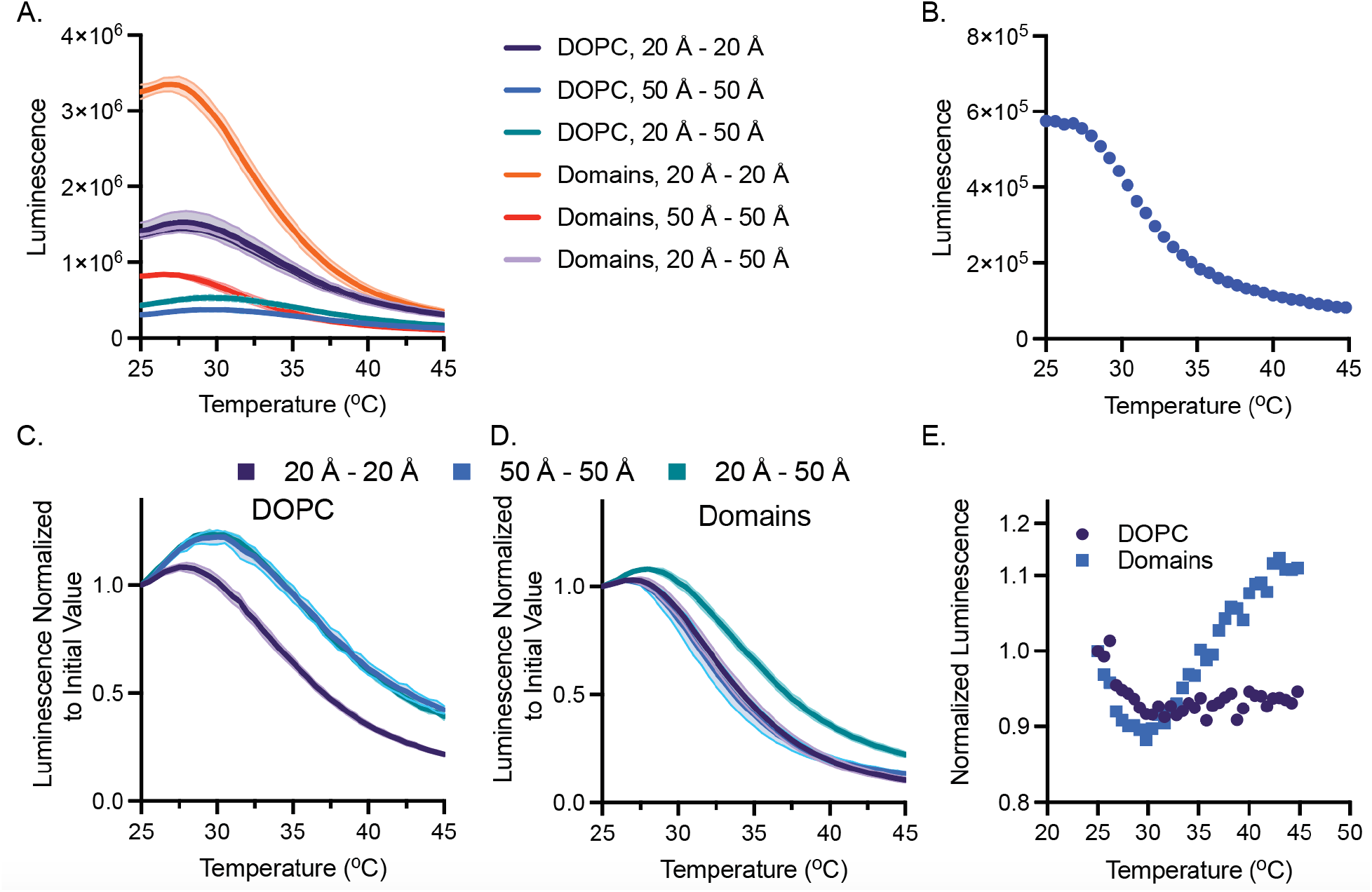
Analysis of split luciferase reconstitution in response to domain dissolution via heating. (**A**) Raw luminescence data of split luciferase constructs in DOPC and domain forming membranes. Luminescence values differ due to differences in expression. Values decrease with increasing temperature, as luciferase is less efficient at elevated temperatures as demonstrated by a luminescence vs temperature for soluble NanoBit (**B**). By normalizing by the initial value, we can compare how luminescence for each combination of proteins changes in (**C**) DOPC and (**D**) domain forming membranes. For domain forming mixtures, transmembrane homodimers decrease more quickly than the heterodimer case. This suggests that homodimers become more dilute as a result of lipid demixing and heterodimers are able to interact more and thus reconstitute luciferase. Data used to generate Fig. 3H. *n=3*, error bars represent the S. E. M. (**E**) An increase in luminescence is also observed in domain forming mixtures when luminescence values are normalized by soluble luciferase luminescence at each temperature.

## References

1. P. Lujan, F. Campelo, Should I stay or should I go? Golgi membrane spatial organization for protein sorting and retention. Archives of Biochemistry and Biophysics. 707, 108921 (2021).

2. Y. Guo, D. W. Sirkis, R. Schekman, Protein Sorting at the trans-Golgi Network. http://dx.doi.org/10.1146/annurev-cellbio-100913-013012. 30, 169–206 (2014).

3. S. Rodriguez-Gallardo, K. Kurokawa, S. Sabido-Bozo, A. Cortes-Gomez, A. Ikeda, V. Zoni, A. Aguilera-Romero, A. M. Perez-Linero, S. Lopez, M. Waga, M. Araki, M. Nakano, H. Riezman, K. Funato, S. Vanni, A. Nakano, M. Muñiz, Ceramide chain length-dependent protein sorting into selective endoplasmic reticulum exit sites. Science Advances. 6, 8237–8248 (2020).

4. R. Prasad, A. Sliwa-gonzalez, Y. Barral, Mapping bilayer thickness in the ER membrane (2020).

5. H. J. Sharpe, T. J. Stevens, S. Munro, A Comprehensive Comparison of Transmembrane Domains Reveals Organelle-Specific Properties. Cell. 142, 158 (2010).

6. J. H. Lorent, K. R. Levental, L. Ganesan, G. Rivera-Longsworth, E. Sezgin, M. Doktorova, E. Lyman, I. Levental, Plasma membranes are asymmetric in lipid unsaturation, packing and protein shape. Nature Chemical Biology 2020 16:6. 16, 644–652 (2020).

7. J. H. Lorent, B. Diaz-Rohrer, X. Lin, K. Spring, A. A. Gorfe, K. R. Levental, I. Levental, Structural determinants and functional consequences of protein affinity for membrane rafts. Nature Communications. 8, 1–10 (2017).

8. T. Harayama, H. Riezman, Understanding the diversity of membrane lipid composition. Nature Reviews Molecular Cell Biology 2018 19:5. 19, 281–296 (2018).

9. K. A. Schwarz, N. M. Daringer, T. B. Dolberg, J. N. Leonard, Rewiring human cellular input–output using modular extracellular sensors. Nature Chemical Biology 2016 13:2. 13, 202–209 (2016).

10. C. Y. Wu, K. T. Roybal, E. M. Puchner, J. Onuffer, W. A. Lim, Remote control of therapeutic T cells through a small moleculeâ€”gated chimeric receptor. Science (1979). 350 (2015),

11. J. G. Rurik, I. Tombácz, A. Yadegari, P. O. Méndez Fernández, S. v. Shewale, L. Li, T. Kimura, O. Y. Soliman, T. E. Papp, Y. K. Tam, B. L. Mui, S. M. Albelda, E. Puré, C. H. June, H. Aghajanian, D. Weissman, H. Parhiz, J. A. Epstein, CAR T cells produced in vivo to treat cardiac injury. Science (1979). 375, 91–96 (2022).

12. Y. Lin, J. Wu, W. Gu, Y. Huang, Z. Tong, L. Huang, J. Tan, Y. Lin, J. Wu, W. Gu, J. Tan, Y. Huang, Z. Tong, L. Huang, Exosome–Liposome Hybrid Nanoparticles Deliver CRISPR/Cas9 System in MSCs. Advanced Science. 5, 1700611 (2018).

13. T. Q. Vu, J. A. Peruzzi, L. E. Sant’Anna, E. W. Roth, N. P. Kamat, Nano Letters, in press, doi:10.1021/ACS.NANOLETT.1C04365.

14. Z. jie Yang, Z. yan Yu, Y. ming Cai, R. rong Du, L. Cai, Engineering of an enhanced synthetic Notch receptor by reducing ligand-independent activation. Communications Biology 2020 3:1. 3, 1–7 (2020).

15. P. Lu, D. Min, F. DiMaio, K. Y. Wei, M. D. Vahey, S. E. Boyken, Z. Chen, J. A. Fallas, G. Ueda, W. Sheffler, V. K. Mulligan, W. Xu, J. U. Bowie, D. Baker, Accurate computational design of multipass transmembrane proteins. Science (1979). 359, 1042–1046 (2018).

16. C. Xu, P. Lu, T. M. Gamal El-Din, X. Y. Pei, M. C. Johnson, A. Uyeda, M. J. Bick, Q. Xu, D. Jiang, H. Bai, G. Reggiano, Y. Hsia, T. J. Brunette, J. Dou, D. Ma, E. M. Lynch, S. E. Boyken, P. S. Huang, L. Stewart, F. DiMaio, J. M. Kollman, B. F. Luisi, T. Matsuura, W. A. Catterall, D. Baker, Computational design of transmembrane pores. Nature 2020 585:7823. 585, 129–134 (2020).

17. F. A. Heberle, M. Doktorova, H. L. Scott, A. D. Skinkle, M. N. Waxham, I. Levental, Direct label-free imaging of nanodomains in biomimetic and biological membranes by cryogenic electron microscopy. Proc Natl Acad Sci U S A. 117, 19943–19952 (2020).

18. M. L. Jacobs, M. A. Boyd, N. P. Kamat, Diblock copolymers enhance folding of a mechanosensitive membrane protein during cell-free expression. Proc Natl Acad Sci U S A. 116, 4031–4036 (2019).

19. G. S. Waldo, B. M. Standish, J. Berendzen, T. C. Terwilliger, Rapid protein-folding assay using green fluorescent protein. Nature Biotechnology 1999 17:7. 17, 691–695 (1999).

20. A. D. Silverman, N. Kelley-Loughnane, J. B. Lucks, M. C. Jewett, Deconstructing Cell-Free Extract Preparation for in Vitro Activation of Transcriptional Genetic Circuitry. ACS Synthetic Biology. 8, 403–414 (2019).

21. C. E. Hilburger, M. L. Jacobs, K. R. Lewis, J. A. Peruzzi, N. P. Kamat, Controlling Secretion in Artificial Cells with a Membrane and Gate. ACS Synthetic Biology. 8 (2019), doi:10.1021/acssynbio.8b00435.

22. E. Sezgin, I. Levental, S. Mayor, C. Eggeling, The mystery of membrane organization: composition, regulation and roles of lipid rafts. Nature Reviews Molecular Cell Biology. 18, 361–374 (2017).

23. H. J. Kaiser, A. Orłowski, T. Róg, T. K. M. Nyholm, W. Chai, T. Feizi, D. Lingwood, I. Vattulainen, K. Simons, Lateral sorting in model membranes by cholesterol-mediated hydrophobic matching. Proc Natl Acad Sci U S A (2011), doi:10.1073/pnas.1103742108.

24. Q. Lin, E. London, Altering hydrophobic sequence lengths shows that hydrophobic mismatch controls affinity for ordered lipid domains (rafts) in the multitransmembrane strand protein perfringolysin O. Journal of Biological Chemistry. 288, 1340–1352 (2013).

25. J. P. Schlebach, P. J. Barrett, C. A. Day, J. H. Kim, A. K. Kenworthy, C. R. Sanders, Topologically Diverse Human Membrane Proteins Partition to Liquid-Disordered Domains in Phase-Separated Lipid Vesicles. Biochemistry. 55, 985–988 (2016).

26. L. v. Schäfer, D. H. de Jong, A. Holt, A. J. Rzepiela, A. H. de Vries, B. Poolman, J. A. Killian, S. J. Marrink, Lipid packing drives the segregation of transmembrane helices into disordered lipid domains in model membranes. Proc Natl Acad Sci U S A. 108, 1343–1348 (2011).

27. D. Lingwood, J. Ries, P. Schwille, K. Simons, Plasma membranes are poised for activation of raft phase coalescence at physiological temperature. Proceedings of the National Academy of Sciences. 105, 10005–10010 (2008).

28. B. Sorre, A. Callan-Jones, J. B. Manneville, P. Nassoy, J. F. Joanny, J. Prost, B. Goud, P. Bassereau, Curvature-driven lipid sorting needs proximity to a demixing point and is aided by proteins. Proceedings of the National Academy of Sciences. 106, 5622–5626 (2009).

29. S. Katira, K. K. Mandadapu, S. Vaikuntanathan, B. Smit, D. Chandler, Pre-transition effects mediate forces of assembly between transmembrane proteins. Elife. 5 (2016), doi:10.7554/eLife.13150.

30. J. Steinkühler, P. Fonda, T. Bhatia, Z. Zhao, F. S. C. Leomil, R. Lipowsky, R. Dimova, Superelasticity of Plasma-and Synthetic Membranes Resulting from Coupling of Membrane Asymmetry, Curvature, and Lipid Sorting. Advanced Science. 8, 2102109 (2021).

31. S. L. Veatch, S. L. Keller, Separation of Liquid Phases in Giant Vesicles of Ternary Mixtures of Phospholipids and Cholesterol. Biophysical Journal. 85, 3074 (2003).

32. J. T. Marinko, J. T. Marinko, A. K. Kenworthy, A. K. Kenworthy, C. R. Sanders, C. R. Sanders, C. R. Sanders, Peripheral myelin protein 22 preferentially partitions into ordered phase membrane domains. Proc Natl Acad Sci U S A. 117, 14168–14177 (2020).

33. A. S. Dixon, M. K. Schwinn, M. P. Hall, K. Zimmerman, P. Otto, T. H. Lubben, B. L. Butler, B. F. Binkowski, T. MacHleidt, T. A. Kirkland, M. G. Wood, C. T. Eggers, L. P. Encell, K. v. Wood, NanoLuc Complementation Reporter Optimized for Accurate Measurement of Protein Interactions in Cells. ACS Chemical Biology. 11, 400–408 (2016).

34. D. H. de Jong, S. Baoukina, H. I. Ingólfsson, S. J. Marrink, Martini straight: Boosting performance using a shorter cutoff and GPUs. Computer Physics Communications. 199, 1–7 (2016).

35. D. H. de Jong, G. Singh, W. F. D. Bennett, C. Arnarez, T. A. Wassenaar, L. v. Schäfer, X. Periole, D. P. Tieleman, S. J. Marrink, Improved parameters for the martini coarse-grained protein force field. Journal of Chemical Theory and Computation. 9, 687–697 (2013).

36. X. Periole, M. Cavalli, S. J. Marrink, M. A. Ceruso, Combining an elastic network with a coarse-grained molecular force field: Structure, dynamics, and intermolecular recognition. Journal of Chemical Theory and Computation. 5, 2531–2543 (2009).

37. T. A. Wassenaar, H. I. Ingólfsson, R. A. Böckmann, D. P. Tieleman, S. J. Marrink, Computational lipidomics with insane: A versatile tool for generating custom membranes for molecular simulations. Journal of Chemical Theory and Computation. 11, 2144–2155 (2015).

38. R. J. Gowers, M. Linke, J. Barnoud, T. J. E. Reddy, M. N. Melo, S. L. Seyler, J. Domanski, D. L. Dotson, S. Buchoux, I. M. Kenney, O. Beckstein, MDAnalysis: A Python Package for the Rapid Analysis of Molecular Dynamics Simulations. Proceedings of the 15th Python in Science Conference, 98–105 (2019).

39. J. A. Peruzzi, M. L. Jacobs, T. Q. Vu, K. S. Wang, N. P. Kamat, Barcoding Biological Reactions with DNA-Functionalized Vesicles. Angewandte Chemie. 131, 18856–18863 (2019).

40. M. Roederer, Spectral Compensation for Flow Cytometry: Visualization Artifacts, Limitations, and Caveats (2001), doi:10.1002/1097-0320.

41. Q. Lin, E. London, Altering hydrophobic sequence lengths shows that hydrophobic mismatch controls affinity for ordered lipid domains (rafts) in the multitransmembrane strand protein perfringolysin O. Journal of Biological Chemistry. 288, 1340–1352 (2013).

